# Human and mouse transcriptome profiling identifies cross-species homology in pulmonary and lymph node mononuclear phagocytes

**DOI:** 10.1101/2020.04.30.070839

**Authors:** Sonia M. Leach, Sophie L. Gibbings, Anita D. Tewari, Shaikh M. Atif, Brian Vestal, Thomas Danhorn, William J. Janssen, Tor D. Wager, Claudia V. Jakubzick

## Abstract

The mononuclear phagocyte (MP) system consists of macrophages, monocytes, and dendritic cells (DCs). MP subtypes play distinct functional roles in steady state and inflammatory conditions. Though murine MPs are well characterized, their pulmonary and lymph node (LN) human homologs remain poorly understood. To address this gap, we created a gene expression compendium across 15 distinct human and 9 distinct murine MPs from lung, LN, blood, and spleen. Human blood MPs and murine spleen MPs served as validation datasets, as the human-mouse MP homologs are relatively well-defined in these tissues. In-depth RNA sequencing identified corresponding human-mouse MP subtypes and determined marker genes shared and divergent across between species counterparts. Unexpectedly, at the gene expression level, only 13-23% of the top 1000 marker genes (i.e., genes not shared across species-specific MP subtypes) overlapped in corresponding human-mouse MP counterparts, indicating a need for caution when translating mouse studies to human gene targets and functions. Lastly, CD88 was useful in both species to distinguish macrophage and tissue monocytes from DCs. Our cross-species gene expression compendium serves as a resource for future translational studies to investigate beforehand whether pursuing specific MP subtypes, or genes will prove fruitful.

## Introduction

Mice are the most commonly used organisms to study human diseases; they breed rapidly and can be genetically modified, accelerating the pace of discovery. A number of similarities with humans suggest that treatments developed in mice may apply to humans. As an example, both species have an immune system that contain mononuclear phagocytes (MPs), granulocytes, innate lymphoid cells, and lymphocytes (Masopust et al., 2017). Immune checkpoint blockades, a Nobel prize-winning discovery, were first observed in the immune system of mice and then translated for therapeutic use in humans (Altmann, 2018); and the development of GM-CSF therapy for pulmonary alveolar proteinosis (PAP) was influenced by observations in mice, where the deficiency of GM-CSF results in PAP (Trapnell et al., 2009).

In spite of these encouraging translational examples, numerous failures in drug development across multiple diseases make it clear that rodent-to-human translation remains an important challenge (Begley and Ellis, 2012). Translational efforts have been challenging across multiple diseases, and there are numerous causes that contribute to these challenges. One key obstacle is the inability to identify functional cellular and molecular homologs in humans and mice. Often the targets identified in mice do not function similarly in humans. Conversely, targets identified in humans are frequently reverse translated into the study of inappropriate murine targets based on naïve assumptions about homology. Therefore, assessing the transcriptome alignment of cross-species human-mouse MPs is crucial.

If equivalence classes of human and murine MP subtypes and their transcriptional markers can be established, studying how murine MPs interact with each other to maintain homeostasis and resolve inflammation could help decipher the mechanisms that go awry in human diseases. Murine MPs have been extensively characterized and demonstrate a clear division of labor during innate and adaptive immunity (Guilliams et al., 2013, Desch et al., 2014, Gibbings et al., 2017, Atif et al., 2015, Kim et al., 2014, McCubbrey et al., 2018, Atif et al., 2018b, Tussiwand et al., 2015). Pulmonary murine MPs consist of alveolar macrophages (AMs), tissue trafficking monocytes, dendritic cell (DCs) subtypes (Tussiwand et al., 2015, Murphy, 2013) and three interstitial macrophages (IMs). Two IMs, IM1 (CD206^hi^MHCII^+^) and IM2 (CD206^hi^MHCII^lo^), display classical macrophage characteristics. The third IM, IM3 (CD206^lo^MHCII^+^), displays macrophage properties but has a higher rate of turnover and expresses pro-inflammatory, monocytic and DC genes (Gibbings et al., 2017, Schyns et al., 2019, Chakarov et al., 2019). Very little is known about IMs in humans, particularly how to identify them, and in both species, humans and mice, how they functionally contribute to homeostasis, inflammation, or disease (Ural et al., 2020, Lim et al., 2018). The importance of knowing more about the transcriptional and functional role of IMs is that they appear to maintain their transcriptional signature across multiple organs, as DCs appear to do (Gibbings et al., 2017, Tamoutounour et al., 2013, Tamoutounour et al., 2012, Epelman et al., 2014, Plantinga et al., 2013).

Circulating monocytes were traditionally viewed as precursors to tissue-resident macrophages. However, we now know that monocytes continuously traffic through non-lymphoid and lymphoid tissue, where they survey the environment. Unless there is a macrophage niche to fill, steady-state monocytes do not differentiate into self-renewing, tissue resident macrophages (Guilliams and Scott, 2017, Bain et al., 2016, Jakubzick et al., 2013, Jakubzick et al., 2017). But during inflammation, monocytes can differentiate into inflammatory and resolving macrophages, which display distinct properties from resident macrophages (Mould et al., 2017, Mould et al., 2019, McCubbrey et al., 2018, Misharin et al., 2017). In lymphoid tissue, even though LN monocytes are highly present in humans and mice, their role in adaptive immunity is less defined than DCs (Jakubzick et al., 2017, Larson et al., 2016, Bosteels et al., 2020).

DCs are potent antigen-presenting cells that link innate and adaptive immunity. In the periphery, DCs can acquire pathogens, traffic through lymphatic vessels to draining LNs, and present exogenous antigens to cognate T cells, precipitating adaptive immunity (Vermaelen et al., 2001, Jakubzick et al., 2006, Jakubzick et al., 2008, Desch et al., 2011, Atif et al., 2018b). In mice, there are two main DC types, Batf3^+^ DC1 and Irf4^+^ DC2. These DCs express distinct transcriptional factors that regulate their development (i.e., which they are named after, among other names), antigen acquisition, processing and presentation capabilities. All in all, although many of the functional cell types and properties of pulmonary MPs have been defined in mice, an assumption made is that similar functional cell types exist in the human lung.

To date, MP cross-species comparisons have been performed for human blood to mouse spleen and human heart to mouse heart (Ingersoll et al., 2010, Bajpai et al., 2018, Bachem et al., 2010, Jongbloed et al., 2010). Although a number of studies have examined the cell surface marker expression and performed single-cell or bulk RNA sequencing (RNA-seq) for human or murine pulmonary MPs (Yu et al., 2015, Yu and Tighe, 2018, Gibbings and Jakubzick, 2018b, Gibbings and Jakubzick, 2018a, Desch et al., 2015, Mould et al., 2017, Reyfman et al., 2018, Bharat et al., 2016, Misharin et al., 2013), a transcriptional alignment of the human-murine pulmonary AMs, IMs, DCs, monocytes, and lymph node (LN) monocytes and DCs is still lacking.

Here we created a gene expression compendium by in-depth RNA-seq to establish the non-diseased transcriptional profile of 15 distinct human and 9 distinct murine MPs, defined by our group and others (Desch et al., 2015, Gibbings and Jakubzick, 2018a, Yu et al., 2015, Yu and Tighe, 2018). The primary focus is pulmonary MPs, especially the understudied LN MPs. However, we also include human blood and murine spleen MPs as control tissues to validate the analytical quality of cross-species pulmonary MP analysis, as homology has been more extensively characterized. Interestingly the overlapping percentage of the top 1000 marker genes defined for the homologous pairs, human blood and murine splenic MPs were similar to those observed for lung and LN MP cross-species counterpart pairs. Overall, we (a) developed a strategy to determine the cross-species counterpart for each MP subtype; (b) compared MP subtypes’ global expression profiles; (c) contrasted specific sets of MP subtype marker genes; and (d) demonstrated how the compendium and subtype analysis can be used to develop new cell surface markers to distinguish macrophage/monocytes versus DC lineages.

## Results

### Identification of human and murine MP populations

We isolated 15 distinct human MPs from four tissue locations: bronchoalveolar space, lung interstitium and draining-LNs from non-diseased donors, and blood from healthy donors. Ten different MP populations were sorted from each lung donor, and 5 different populations were sorted from each blood donor (Figure 1A and Figure S1). Putative MP subtype labels were assigned based on 1) analyses of over 100 human lungs examined in our laboratory for given MP populations consistently observed regardless of age and gender and 2) well-established cell surface markers considered to selectively identify macrophages or DCs (Figures 1A, 1C, S1) (Gibbings and Jakubzick, 2018b, Gibbings and Jakubzick, 2018a, Desch et al., 2015, Bharat et al., 2015, Misharin et al., 2013, Yu et al., 2015, Yu and Tighe, 2018, Guilliams et al., 2016, Bharat et al., 2016). For each human lung, we isolated seven MP populations: four distinct macrophage populations (AM, IM CD36, IM HLADR, and IM CD1c^+^), two CD1c^+^CD36^-^ DC subsets (DC CD206^-^ and DC CD206^+^), and CD14^+^ monocytes. Peribronchial LNs contributed CD14^+^ LN monocytes and two LN DC populations (DC CD1a^hi^ and DC CD1a^lo^). Last, we sorted well-defined blood (BL) MPs including two monocyte populations (Mo CD14^+^CD16^-^ and Mo CD14^+^CD16^+^) and three classical CD11c^+^HLADR^+^ DCs (DC Clec9a^+^, DC CD1c^+^, and DC CD1c^-^) (Desch et al., 2015, Ingersoll et al., 2010, Huysamen et al., 2008). In total, we sorted and analyzed 66 human MP samples by RNA-seq, with three-to-six replicates per MP subtype.

**Figure 1.**
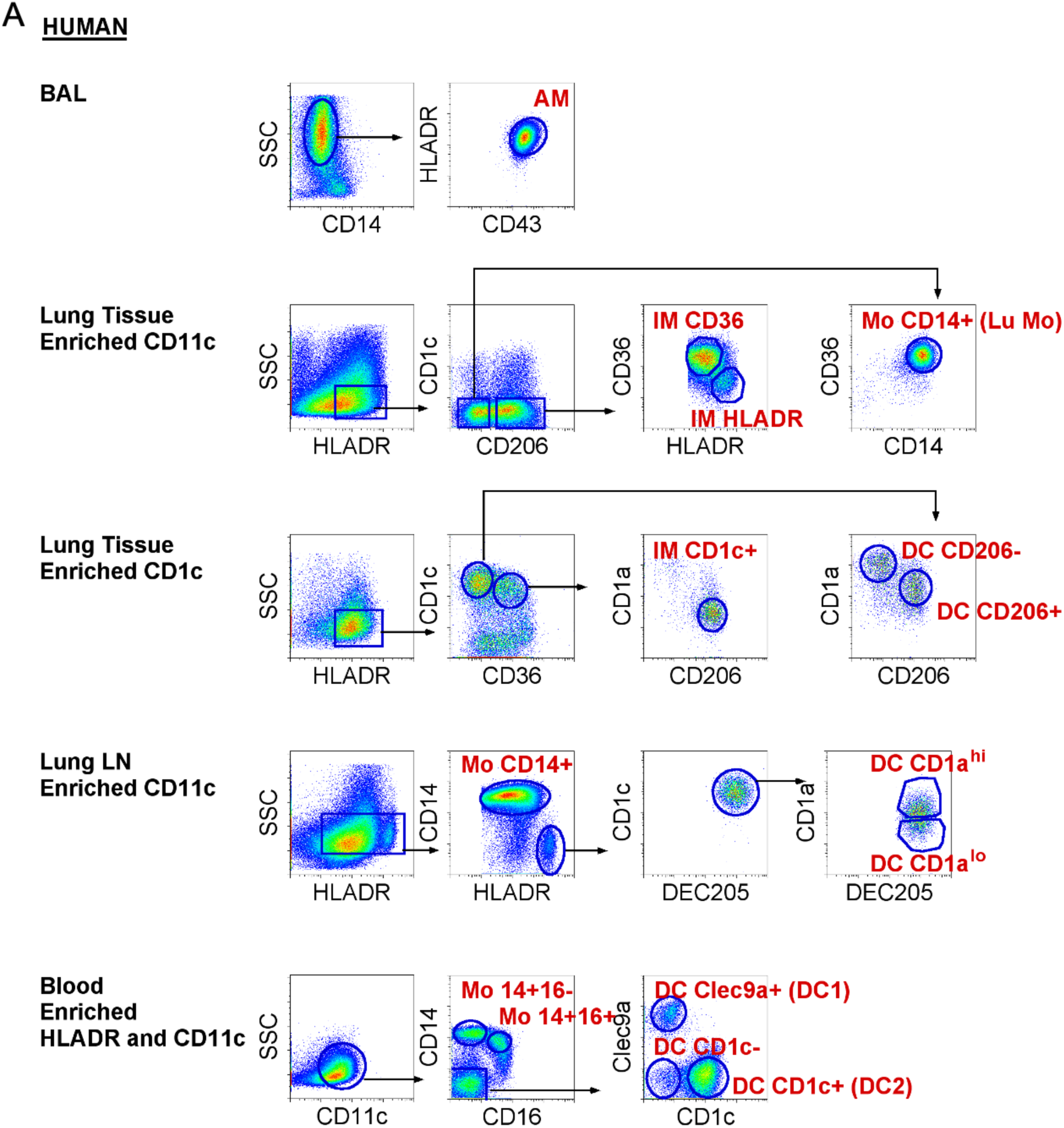
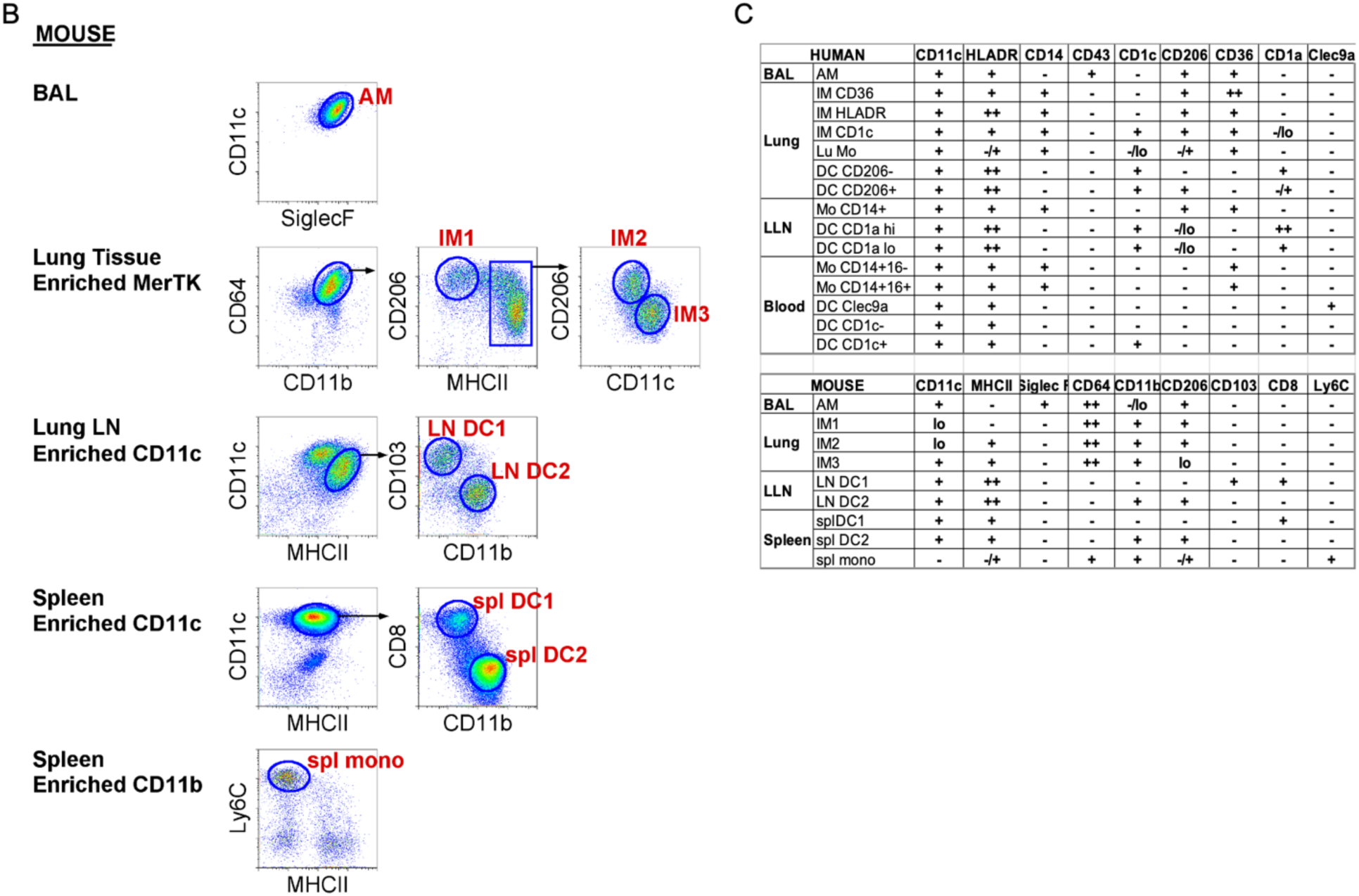
Gating strategy to sort human and mouse mononuclear phagocytes for RNA isolation A) Cell suspensions were magnetically enriched, as indicated, from bronchoalveolar lavage (BAL), lung digest, lung-draining lymph nodes (LLN), or blood (BL). Fifteen different populations of mononuclear phagocytes (MPs, labeled in red) were FACS sorted, as shown, after gating to select Live; Single; CD45^+^; Lineage^-^ (Lin) cells. Side scatter properties (SSC) were used to separate AMs-SSC^hi^ from other MPs-SSC^int^. BAL AMs were identified by co-expression of CD43 and HLA-DR and lack of CD14. Pulmonary Interstitial macrophages (IM) were defined as CD206^+^ and CD14^+^ and separated into three populations based on their differential expression of CD36, HLA- DR and CD1c. Pulmonary monocytes were taken from the CD206^-^ fraction (Lu Mo). In lung and LLN, dendritic cells (DC) were characterized as CD14^-^, CD36^-^ cells that express CD1c and were separated into two populations in each organ based on the differential expression of CD206 and CD1a. In the LLN, DEC205 was also used to define DCs. Similarly, in blood CD14 and CD16 expression was used to separate monocytes (Mo) from DCs. BL DC subpopulations were defined on the basis of Clec9a or CD1c expression. **B)** Cell suspensions were magnetically enriched as indicated from bronchoalveolar lavage (BAL), lung digest, lung-draining lymph nodes (LLN), or spleen. Nine different populations of mononuclear phagocytes (MPs, labeled in red) were FACS sorted as shown, after gating to select Live; Single; CD45^+^; Lineage^-^ (Lin) cells. Alveolar macrophages (AM) co-express CD11c and Siglec F in the BAL. In the lung, interstitial macrophages (IM) are defined by expression of MerTK, CD64 and CD11b, and can be divided into three subpopulations based on the differential expression of CD206, MHCII and CD11c. In the LLN and spleen DCs co-express CD11c and MHCII separated by differential expression of either CD103 or CD8 (DC1) and CD11b (DC2). Monocytes are distinguished from other splenic MPs by expression of CD11b and Ly6C. **C)** Table illustrating MP subsets cell surface marker expression.

In the mouse, we isolated nine distinct murine MPs from four tissue locations: bronchoalveolar space, lung tissue and draining-LNs, and spleen (Figure 1B and Figure S2). We isolated four lung macrophage populations (AM, IM1, IM2, and IM3) and two LN migratory DCs (CD103^+^ DC1 and CD11b^+^ DC2), distinguished from lymphoid resident DCs by high MHCII expression (Figure 1B). In addition, two splenic DC populations (CD8^+^ DC1 and CD11b^+^CD4^+^ DC2) and Ly6C^hi^ monocytes were sorted. Since mice lack detectable DCs in circulation, splenic DCs were included in this study as the comparators to human blood DCs (Haniffa et al., 2012, Bachem et al., 2010, Schlitzer et al., 2013, Reynolds and Haniffa, 2015, Ingersoll et al., 2010). The spleen lacks afferent lymphatics exclusively representing a source of resident, lymphoid DCs, in contrast to tissue experienced, migratory DCs. The total number of murine MP samples sorted and analyzed by RNA-seq was 27, with three replicates per MP subtype.

### Confirming human MP subtypes based on defined transcriptional profiles

To validate our designations of sorted human MPs as either macrophages, monocytes or DCs, we examined the expression of classical macrophage/monocyte and DC genes (Figure 2) (Scott et al., 2014, Wu et al., 2016, Satpathy et al., 2012). As anticipated, most of the macrophage/monocyte genes such as *MAFB*, Fc gamma receptors, lysosomal proteases and associated membrane proteins were most strongly expressed in human MP subtypes putatively labeled as macrophages or monocytes (Figure 2) (Tussiwand et al., 2012, Satpathy et al., 2012, Wu et al., 2016). Conversely, classical DC genes such as *HLA-DR*, *CD1C, CCR7*, *CD40*, *CD80*, *LAMP3*, *IRF4*, *IRF8*, *FLT3* and *ZBTB46* were strongly expressed in human pulmonary and LN MPs, putatively labeled as DCs (Figure 2) (Wu et al., 2016, Satpathy et al., 2012). In fact, pulmonary and LN DCs showed higher relative expression of classical DC genes than did blood DCs. For example, Figure 2 shows high expression of *CCR7*, a key DC signature gene, in pulmonary and LN DCs. DCs require CCR7 for both migration from tissue to LNs via afferent lymphatics and T cell zone localization (Forster et al., 1999). Overall, based on cell surface markers and transcriptional data, we were able to confirm the appropriate labeling of human pulmonary and LN MPs.

**Figure 2.**
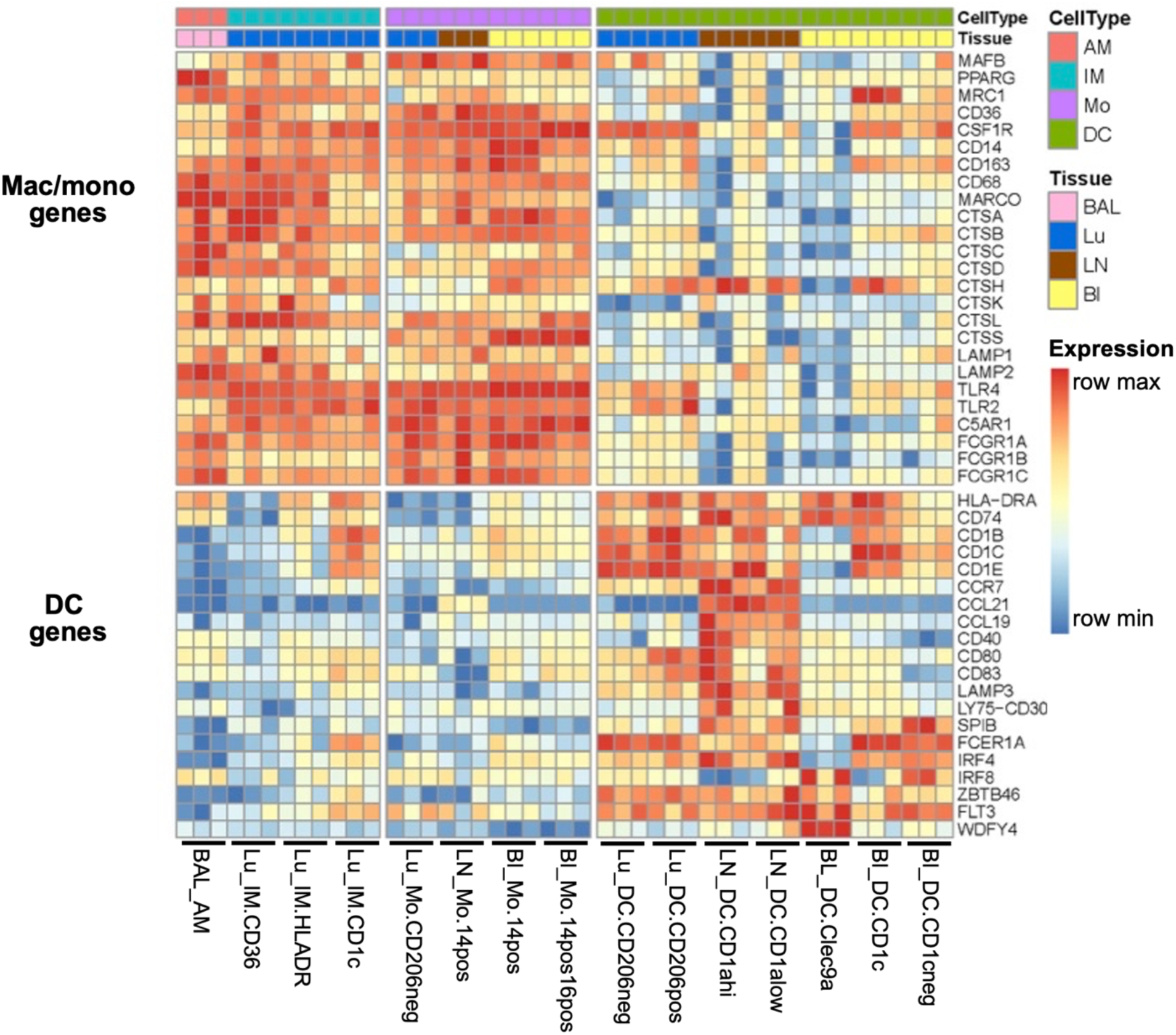
Confirmation of human MP subtypes based on classical gene signatures. Expression of signature DC and macrophage transcripts from RNA-seq are shown as normalized values. Data first undergo a variance stabilizing transformation on read counts, where the donor identity has been regressed out to mitigate any donor effect. The data for each gene is then min-max scaled into the range [0,1] to emphasize which MP subtypes have the minimum or maximum expression for a given gene. Though the number of replicates ranges from three to six, only the first three replicates are shown for each subtype.

### Combined human and murine MP data grouped subtypes by AM, IM, mono or DC lineage

A cross-species data set of RNA-seq reads was obtained for all autosomal genes with a 1:1 homology between human and mouse. Sex chromosome genes were omitted to avoid variation based on donor sex. To illustrate the relative correspondence between human and murine MP subtypes, we aligned human and murine MP expression data, borrowing methods originally developed for single-cell RNA sequencing (scRNA-seq) to account for species-specific effects on gene expression (Butler et al., 2018). Canonical correlation analysis (CCA), followed by alignment and t-Distributed Stochastic Neighbor Embedding (t-SNE) were applied to quantile-normalized human and murine MP data (Figure S3-S4) (Bolstad et al., 2003). The unbiased analysis used 885 genes shared between human and mouse from among the top 2000 most highly variable autosomal genes in either species, capturing similarities among global gene expression profiles without knowledge of MP subtype identity.

The combined human-mouse t-SNE plots illustrate that clusters of cells with similar transcriptional profiles are organized by MP subtype rather than species or tissue type. Cell populations with similar profiles included a mix of human and mouse cells and multiple tissues of origin, indicating that species and tissue type were not the main sources of variation in gene expression (Figure 3A). Crucially, the transcriptional profiles were instead clustered by MP cell type, with AMs, monocytes, and DCs forming tight clusters, followed by IMs (Figure 3A). To facilitate comparisons, samples were further grouped into six groups, labeled with an indication of their lineage (Figure 3B). Group-1 AM contains AMs from BAL (macrophage that is tissue-specific). The IM (Group-2 IM) group separates from AMs, as previously observed in mice (Gibbings et al., 2017). Group-3 (Mono) is a pure cluster of monocytes across species, sampled from lung, blood and spleen. Despite the fact that AM and IM share expression of a number of macrophage core signature genes, their overall gene expression is strongly influenced by their local environment, alveolus versus interstitium (Gautier et al., 2012, Gibbings et al., 2017). Group-2 (IMs) also includes CD14^+^ monocytes from the LN. Group-2 IMs also includes CD14^+^ monocytes from the LN. This likely reflects their shared origin as monocyte-derived populations that matured in the lung environment.

**Figure 3.**
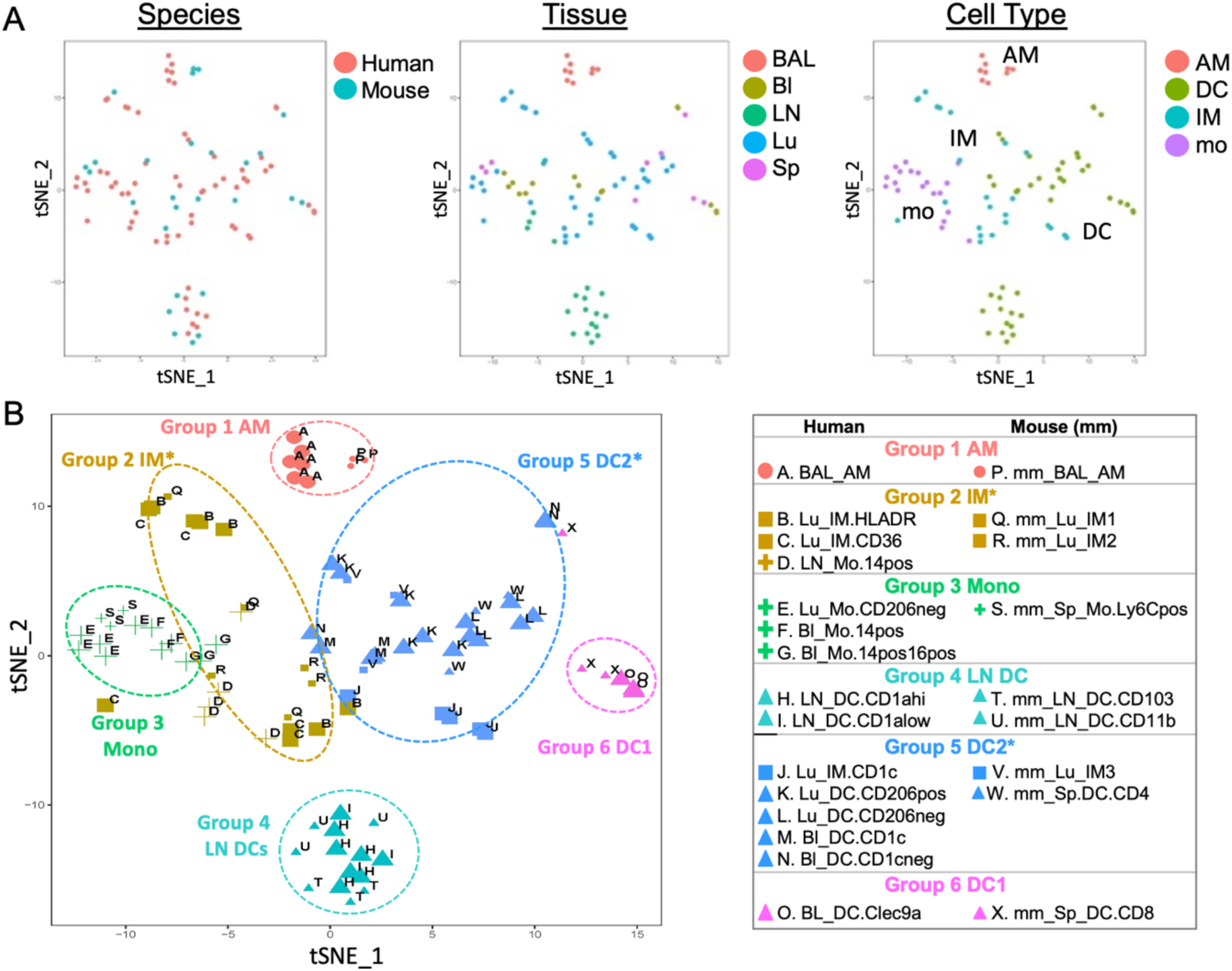
Visualization of MP subtypes across species. Quantile-normalized VST subject-regressed data from 3-6 donors per subtype for 15 human and 9 murine MP subtypes was subjected to the Seurat v2.0 pipeline to visualize human and mouse data within a common two-dimensional space. A) Samples in the resulting 2-D projection are colored by species, tissue, or cell type. B) Samples are grouped into six clusters based on later shared marker gene analysis (Figure 4). Shape indicates cell type, size indicates species, and color indicates group assignment. MP subtypes are denoted by the letters A through X. Note each group contains related subtypes across species.

The DCs are distributed across three distinct groups: Group-4 LN DCs, Group-5 DC2 and Group-6 DC1 (Figure 3B). LN DCs in Group-4 form a distinct cluster, clearly separated from the other DCs, suggesting a maturation or functional program distinct from blood or splenic DCs. Group-5 DC2 contains the human lung/blood DCs and murine splenic CD4^+^ DCs, suggesting an overall DC2 (Irf4-dependent) lineage. Surprisingly, Group-5 also contains human lung CD1c^+^ IMs and murine lung IM3s. These subtypes were isolated using classical IM cell surface markers, such as CD64, CD36, and MerTK, yet comparison of their global expression patterns suggests this IM subtype in human and mouse also exhibit some DC expression programs. This draws a parallel to the Langerhans cell in skin, that has both macrophage and DC-like functions (i.e., a self-renewing macrophage that migrates to LN and presents antigen), and expression of macrophage and DC-specific transcription factors, *mafb* and *zbtb46* (Wu et al., 2016). Lastly, Group-6 DC1 contains human blood Clec9a^+^ DCs and murine splenic CD8^+^ DCs, the expected human-to-mouse counterparts of the Batf3-dependent DC1 lineage (Hildner et al., 2008). Overall, the unbiased computational analysis of global gene expression data demonstrates that across species the majority of MP subtypes group by cell type —AM, IM, monocyte or DC lineage— across species.

### Marker gene analysis identifies the closest cross-species homologue per MP subtype

We develop a novel, robust strategy to determine the maximally corresponding MP subtype between human and mouse. Other methods to establish homologous MP populations have relied on global correlation (Travaglini et. al. bioRxiv https://doi.org/10.1101/742320) or marker gene set enrichment (Chakarov et al., 2019), which can be affected by the set of genes used in the calculations. For instance, genes with low expression levels and housekeeping genes can bias global correlations so that cell types appear similar across many cell types, failing to discriminate (Figure S5A). Strong correlations across multiple cell types can occur even when using only highly variable genes and may disagree with the global correlations (Figure S5B). Marker gene sets often overlap for closely related cell types, especially when not accounting for the level of uniqueness of expression for that cell type. Thus, enrichment analysis based on intersections of unranked marker gene lists can obscure the dominant functional homolog (Figure S6A). If the closest homolog is chosen as the cell type with the largest overlapping marker set, the closest homolog using the top 50 marker genes may be different than the closest homolog using the top 1000 (Figure S6B). Ideally, the best homolog would be the cell type that continues to have the largest overlap, even as the size of the (ordered) marker set increases. Our approach captures this intuition to determine the maximally corresponding MP subtype between species.

First, our approach prioritizes potential marker genes for each MP subtype within each species. Marker genes should be highly and preferentially expressed within a single subtype over all other subtypes within a species. In total, 15 marker gene sets for human and 9 marker gene sets for mouse are evaluated from a dataset consisting of 11,959 genes across 93 samples. Each gene is ranked for each subtype by a score that multiplies two factors: 1) the level of expression in a given subtype relative to the median expression over all subtypes; and 2) how differentially expressed the gene is in that subtype over all other subtypes (Rosenthal, 1978). Thus, genes with scores earlier in the ranking for a subtype are both most highly expressed and most differentially expressed for that subtype.

To achieve a high score, marker genes for each of the 15 human and 9 murine subtypes must be differentially expressed with respect to each remaining subtype within their species, in contrast to other marker gene identification methods that consider the subtype versus the union of all samples from the other subtypes. Note that as a result, core signature genes at the level of macrophages, monocytes or DCs will not be considered highly ranked, since those genes are shared among the multiple related subtypes, and thus receive a low score for differential expression. In this way, we take advantage of the granularity at which the subtypes were isolated to find the maximally corresponding homologue based on marker genes essentially unique to that subtype.

Once marker genes are ranked, we can examine the degree of overlap of marker sets between species to determine the closest cross-species match. Our method takes into account the level of uniqueness of expression for that cell type and prioritizes cell types with strong overlap regardless of the size of the marker set considered. We develop a method based on Correspondence-at-the-Top (CAT) plots, previously used to compare expression results across experimental platforms (Irizarry et al., 2005). Where a single Venn diagram can depict the intersection (i.e., percentage overlap) of two unordered and fixed sized lists, CAT plots visualize the intersection percentage of two ranked lists as the list size progressively increases (Figure 4A). Using an MP subtype in one species as the reference, each MP subtype in the opposing species is compared to the reference to determine the percent overlap as the marker gene set size increases, i.e., comparing the top-5 genes, the top-10 genes, etc (examples illustrated in Figure 4A, all human and mouse MPs in Figure S7-S8). As seen in the CAT plots, generally the curves stabilize and one cell type dominates as the best cross-species match within the top 1000 genes (vertical dashed line).

**Figure 4.**
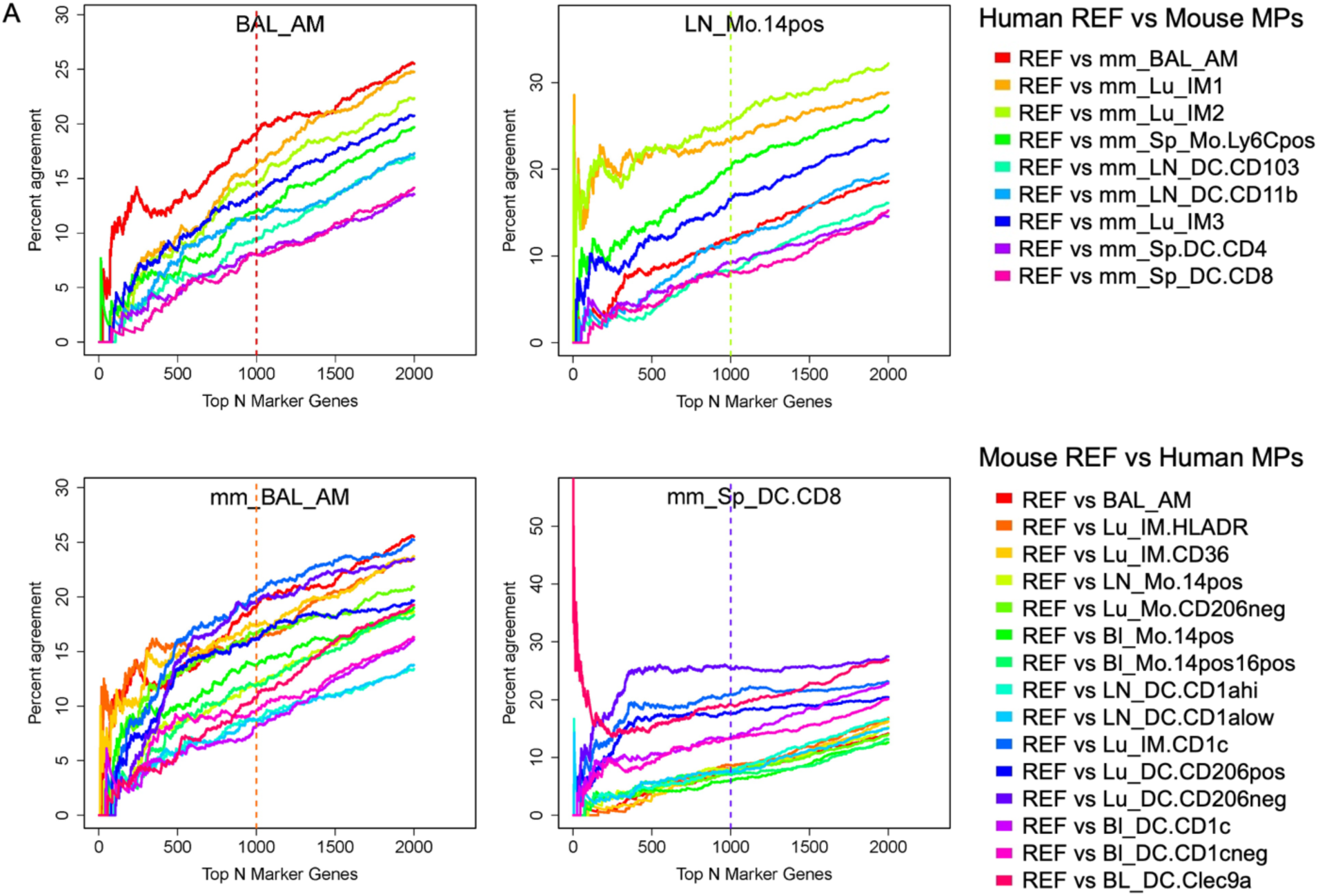
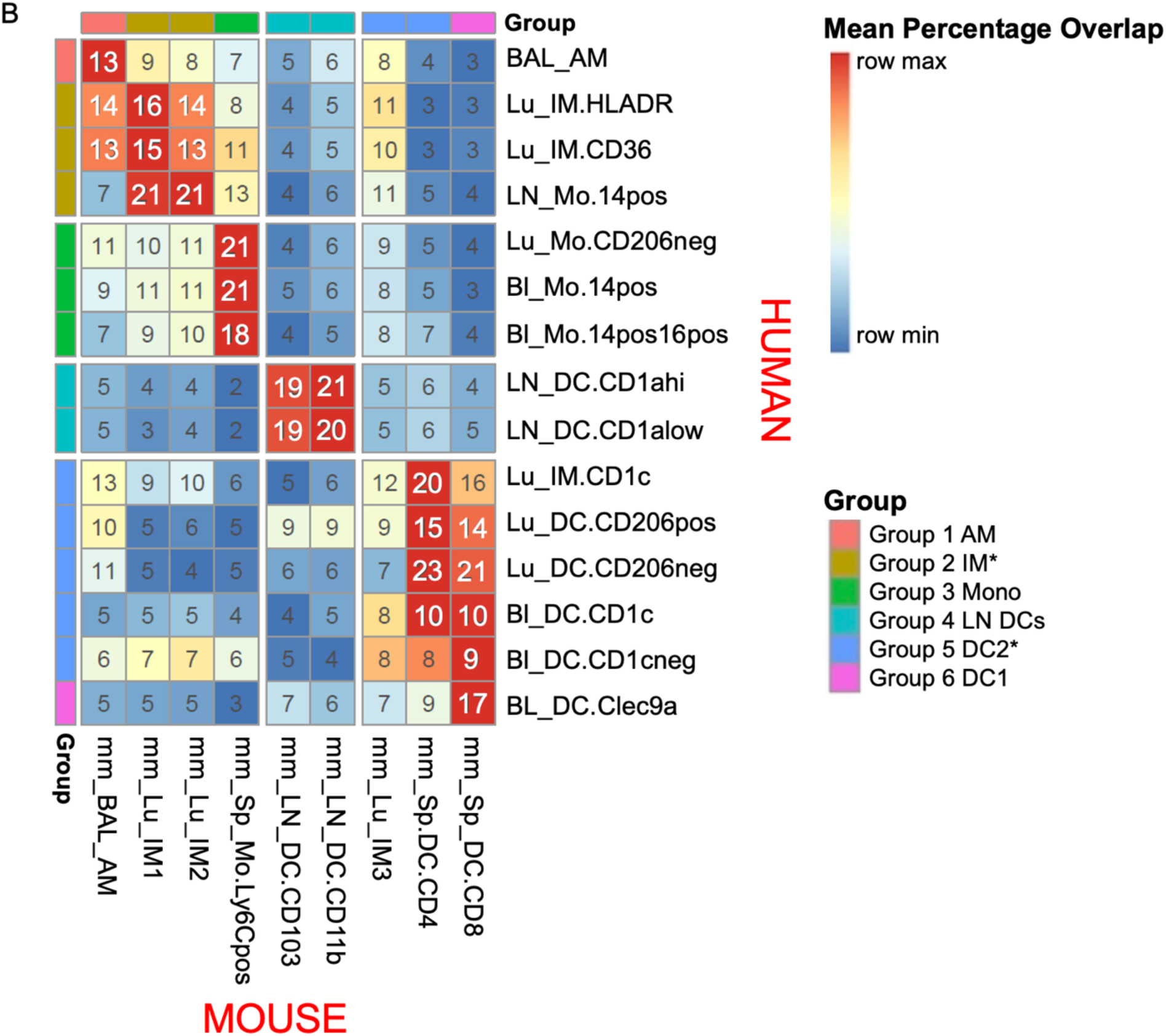
Correspondence of MPs across species. Candidate marker genes were ranked by their score, which multiplies their expression levels, scaled relative to the median expression, by their marker gene Z-score (see Methods). Top ranking marker genes, the most important marker gene, are on the left, with higher ranks (up to 2000) on the right. Percentage agreement is the percentage of genes identified as critical markers in the reference species that are also identified as critical markers for the comparison species up to that rank. For example, at rank 1, if the top-ranked marker for human is also the top-ranked for mouse, percentage agreement would be 100%, and otherwise it is 0%. At rank 100 on the x-axis, a value of 10% would indicate that of the marker genes in the top 100 in the reference species, 10% of them are in the top 100 of the comparison species. Each line shows a different cell population in the comparator species. For example, for the top left panel A, human BAL_AM is the reference cell type, and murine (mm) is the closest match across all rankings, which can be seen because it has the highest percentage overlap (highest on y axis). **A)** For each reference subtype the top panels show humans as the reference species, and the bottom panels show mouse as the reference species (signified by mm). The proportion of agreement of ranked marker gene lists versus ranked marker gene lists for each candidate subtype in the other species was calculated for progressively larger list sizes and visualized with a Correspondence At The Top (CAT) plot for a select set of MP subtypes. The color of the vertical line indicates which subtype had the highest mean CAT overlap across all list sizes from 1 to 1000 with respect to the reference. **B)** The mean CAT overlap over list sizes 1 to 1000 is shown for each subtype pair. Rows are scaled by the max and the min to emphasize the highest subtype correspondence for the human subtypes. Rows and columns are ordered to cluster similar subtypes. Clusters of MP subtypes are then annotated by group number and these groupings are used for the coloring in Figure 3B.

One option for choosing the closest match between species would be the cell type with the maximum match at the top 1000 genes. However, marker sets are ordered by degree of unique expression for the cell type, so maximizing at any one list size would not distinguish between a pair that agrees quickly in the ranking and maintains agreement up to that list size (i.e., strong, robust agreement among the most unique marker genes) from a pair that achieves high agreement only later in the ranking (i.e., includes mainly marker genes appearing in multiple cell types or more lowly expressed genes). To be robust to marker set size and determine which subtype quickly and consistently dominates, we instead choose to summarize the correspondence of each subtype pair across species as the mean percentage overlap of the corresponding marker gene lists for the pair across all top list sizes from 1 to 1000, referred to henceforth as mean CAT overlap (Figure 4B). Overall, the mean CAT overlap ranges between 2% and 23%. The closest cross-species homologue for a given MP subtype is then chosen as that subtype in the opposing species with the highest mean CAT overlap, which ranges from 13% to 23%. A potential factor contributing to the low percentages is the fact that our ranked marker genes are selected to be those expressed exclusively in a single cell type. Core signature genes appearing in multiple cell types are discarded. Including these would substantially increase the overall percentage overlap but would be less meaningful because these genes are not diagnostic of one cell type. Of note, the clustering of MP subtypes seen in the previous t-SNE plot analysis of global expression is echoed in the CAT plot analysis of marker gene sets (Figure 3B, Figure 4B). This suggests that despite the low overlap by our strict marker gene definition, the same biological correspondences are found as in the unbiased analysis.

### Human MPs correspond to expected mouse MPs except CD1c^+^ IMs and LN monocytes

Most of the human MPs align best with a single murine cell type. For example, AMs correspond across species, with a 13% mean CAT overlap, especially early in the ranking among the top 200 marker genes (Figure 4A and 4B, Group-1 AM). Human blood and lung monocytes highly corresponded directly to murine splenic monocytes with 21% mean CAT overlap (Group-3 Mono). Human blood Clec9a^+^ DCs are most similar to murine splenic CD8^+^ DCs, with 17% mean CAT overlap (Group-6 DC1); unexpectedly, these Clec9a^+^ DCs are not strongly aligned with murine LN CD103^+^ DCs (7% mean CAT overlap), as would be hypothesized based upon the shared Batf3-dependent lineage of CD8^+^ splenic DCs and CD103^+^ LN DCs in the mouse. Instead LN DCs are highly correspondent across species (Group-4 LN DCs) suggesting that DC maturation state or tissue imprinting may have a greater effect on gene expression.

On the other hand, there are a few human MPs that share homology with more than one mouse MP subtype. Human IM HLADR and IM CD36 are most similar to murine IM1 and IM2, with 13-16% mean CAT overlap (Figure S7 and Figure 4B, Group-2 IM), demonstrating correspondence across IMs, as expected. However, human IM CD1c is most similar to murine splenic CD4^+^ DC (20% mean CAT overlap) and less similar to its expected counterpart murine IM3 (12% mean CAT overlap) (Figure S7 and Figure 4B, Group-5 DC2). Note, however, inversely the murine IM3 is most similar to human IM CD1c, as anticipated (Figure S8 and Figure 4B).

In the LN, human LN monocytes correspond highly with murine IM1 and IM2 (21% mean CAT overlap) instead of splenic monocytes (13% CAT mean overlap, 3^rd^ highest) (Figure 4A and Figure 4B, Group-2 IM). This was also evident in the t-SNE plots (Figure 3B), reflecting the shared characteristics of differentiated monocytes in tissue.

### Murine MPs match expected human MPs, yet exhibit evidence of dual expression programs

We also assessed the inverse correspondence, taking each mouse MP and identifying the most similar human MP (Figure S8 and Figure 4). Murine AMs (Figure 4B, Group-1 AM) are highly similar to human AMs (13% mean CAT overlap), especially when comparing up to the top 200 genes; thus, sharing the most unique AM marker genes (Figure 4A). However, among the top 200 to 1000 genes, human IM CD1c and IM HLADR (Group-2 IM) begin to overlap increasingly (13-14% mean CAT overlap), suggesting murine AMs share dual characteristics with human AMs and IMs. Murine LN DCs and splenic monocytes were most similar to human LN DCs and lung/blood monocytes, respectively (Group-4 LN DCs ∼20% mean CAT overlap and Group-3 mono 21% mean CAT overlap), as anticipated.

Interestingly, where human blood Clec9a^+^ DCs align most strongly with their expected counterpart mouse splenic CD8^+^ DCs, the inverse is slightly different depending on the number of top genes analyzed (Figure 4, Group-6 DC1). Averaging the percent overlap over lists of size 1 to 1000, murine CD8^+^ DCs correlate best with the human lung CD206^-^ DCs (Group-5 DC2, 21% mean CAT overlap) as opposed to Clec9a^+^ DCs (17% mean CAT overlap). Similar to the results for murine AMs, murine splenic CD8^+^ DCs do match their expected homologue, Clec9a^+^ DCs, best when considering only the first top 1 to 50 ranked genes (Figure 4A). This suggests that the transcriptional program of mouse splenic DCs shares similarities with both human blood Clec9a DCs and lung CD206-DCs.

Murine IM1 and IM2 correspond best to human LN monocytes, with 21% mean CAT overlap, followed by human HLADR and CD36 IMs, with ∼15% mean CAT overlap (Figure 4B Group-2 IM and Figure S8). Murine IM3 corresponds highly with human LN monocytes as well as IM HLADR (11% mean CAT overlap) (Figure 4B Group-2 IM and Figure S8), though IM3 matches most strongly with human IM CD1c (12% mean CAT overlap) (Figure 4B Group-5 DC2).

Overall, CAT plot analysis of marker gene sets highlights cross-species correspondences and defines the best overall counterpart for human and murine MPs within our MP subtypes (Figure 4B). In cases where the predicted counterparts do not match expectations, the analysis gave evidence that the marker genes of the expected homologue did appear early in the ranking (i.e., among the top-50 marker genes) suggesting multiple MP expression programs exist in some subtypes when patterns across a larger set of top marker genes (i.e., 1000) are analyzed.

### Conserved and distinct genes expressed in well-defined human-mouse homologs

Expression data for the top marker gene sets are first examined in three well-defined human-mouse homologs: AMs, monocytes and DC1 (Figure 5). The functional properties of genes mentioned below are outlined in Table SI. Expression of the top 50 marker genes in the reference MP subtype is shown in the opposing species (Figure 5-7, Figure S10-S33). Note that the best match suggested visually among expression of just the top 50 genes may differ from the assigned best match maximizing mean CAT overlap computed from the top 1000 genes, thus motivating the decision to highlight the top three closest subtypes of the opposing species.

**Figure 5.**
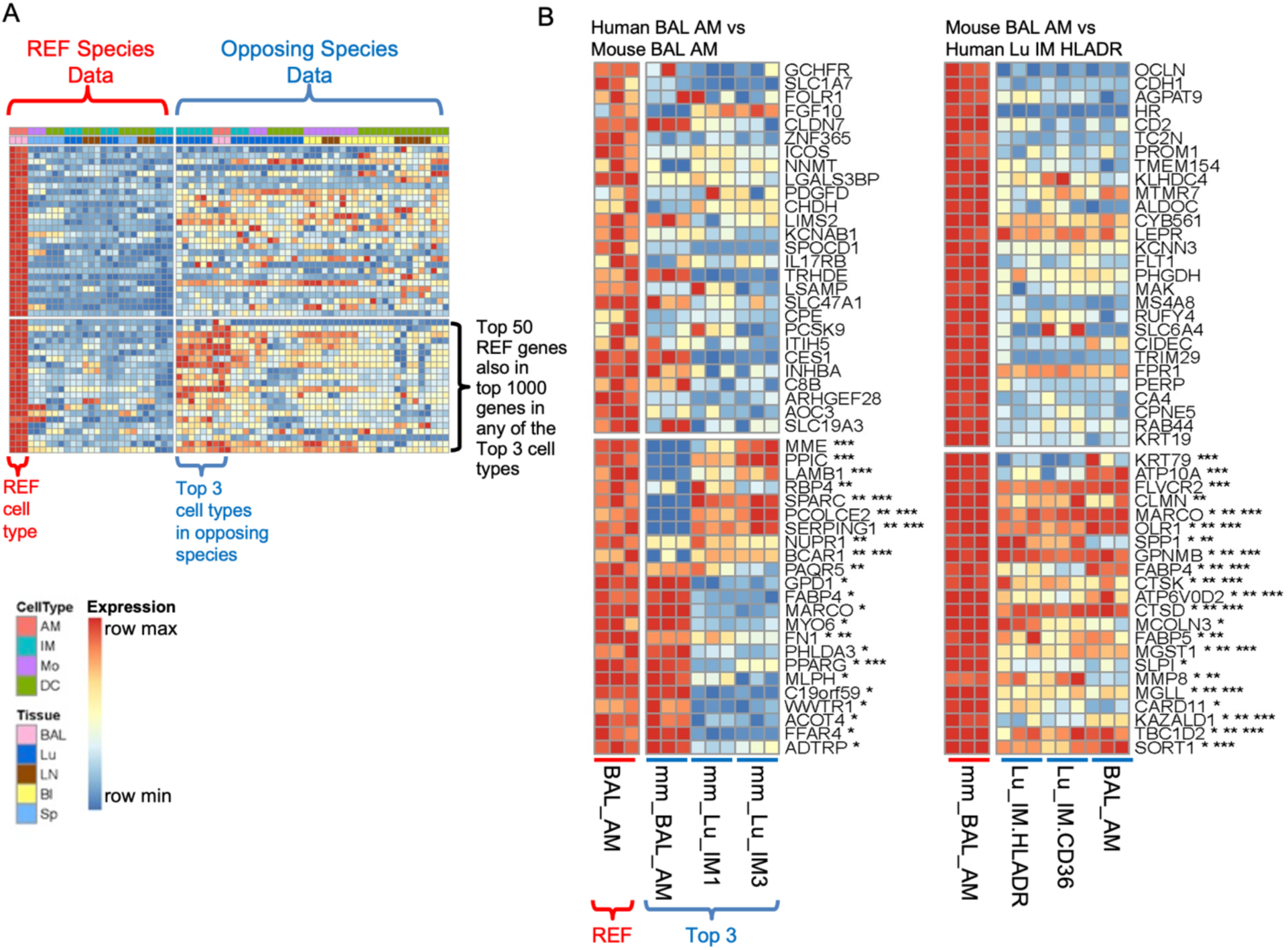
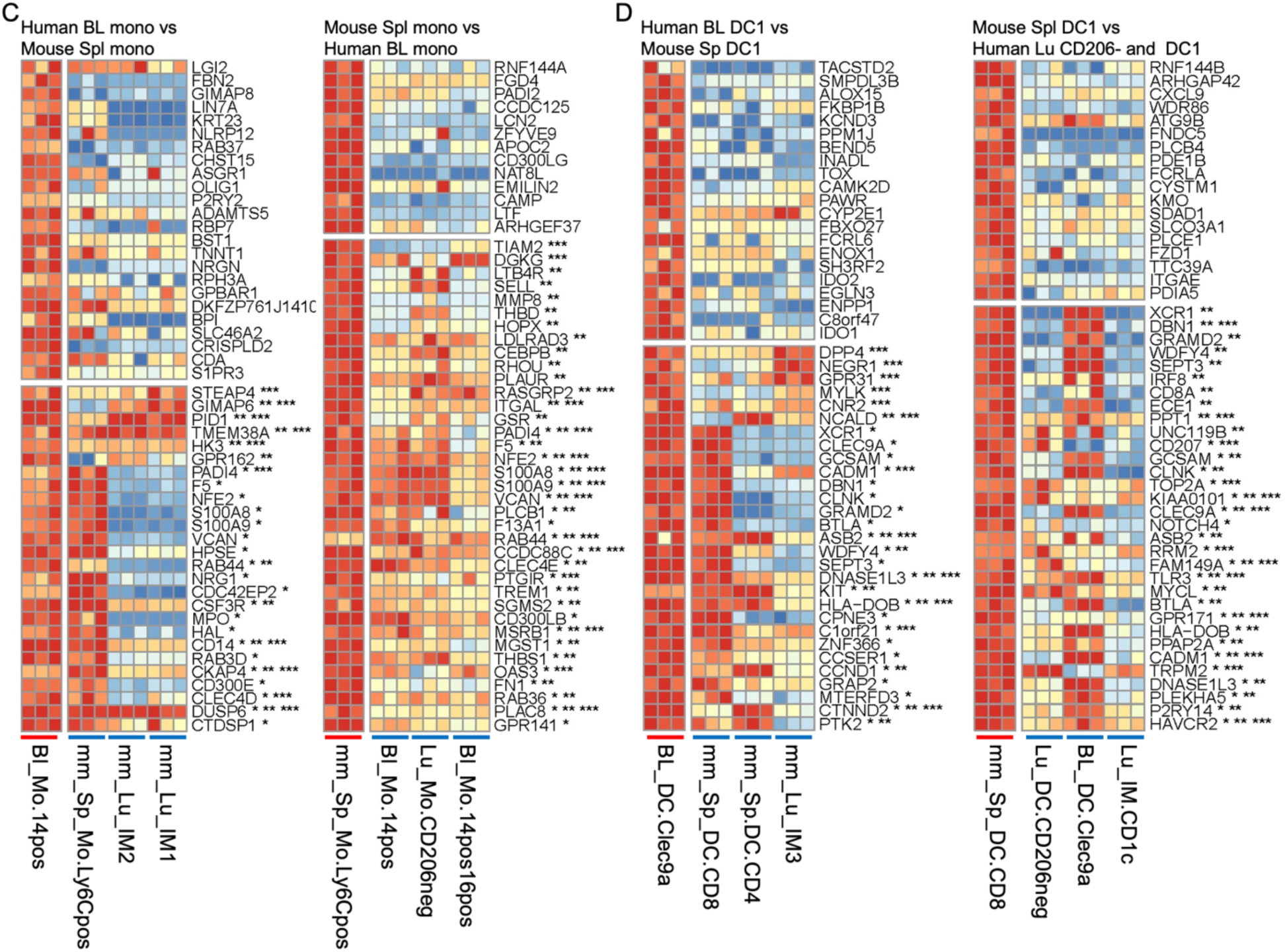
Well-defined human-mouse homologs. The expression values are shown for the top 50 marker genes in the reference cell type. The cell types are ordered by degree of overlap with the reference cell type, if within the same species, or by the mean CAT overlap shown in Figure 4B if in the other species. The data are scaled across the row by the gene min and max within each species to emphasize in which cell types the gene is most expressed. Genes among the top 50 in the reference that also appear in the top 1000 ranked genes in the 1^st^, 2^nd^ or 3^rd^ best match marker list appear towards the bottom and are annotated with *, **, or *** respectively. Note that annotation by an asterisk does not mean that the gene must also appear in the top 50 marker genes in the other species, only that it appears in the top 1000 genes for the top 3 matches. Also, visualizing the top 3 matches rather than just the best match can help reveal cases when the best match, as determined using 1000 reference marker genes, may differ from the best match suggested by comparing expression of just the top 50 reference genes, in which case the CAT plots can inform the reason for the perceived discrepancy (Figure S7-S8). **A)** Example depiction of all data in all samples for a given reference and the top 3 closest matches according to Figure 4B. **B)** Highest correspondence for human and mouse AMs. Three replicates for the reference and three closest subtypes of the opposing species are shown. Data with all replicates can be found in Figure S10-S33 and is useful to consult for additional information about specific genes or cell types. **C)** Reciprocal highest correspondence for human Bl and mouse Spl monocytes. Three replicates for the reference and three closest subtypes are shown. **D)** Highest correspondence for human Bl DC Clec9a^+^ and mouse Spl CD8^+^ DC1. Three replicates for the reference and three closest subtypes are shown.

**Figure 6.**
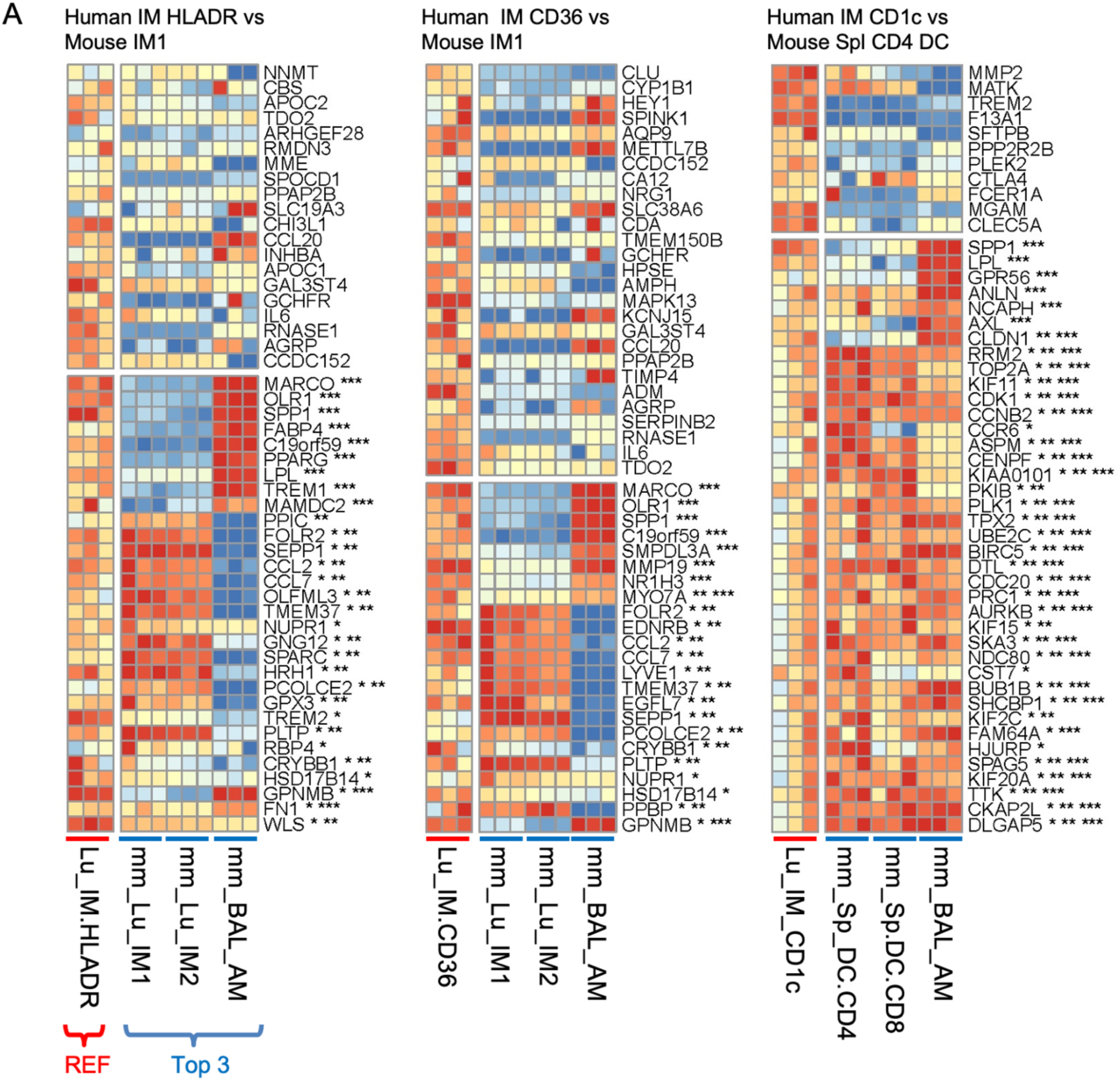
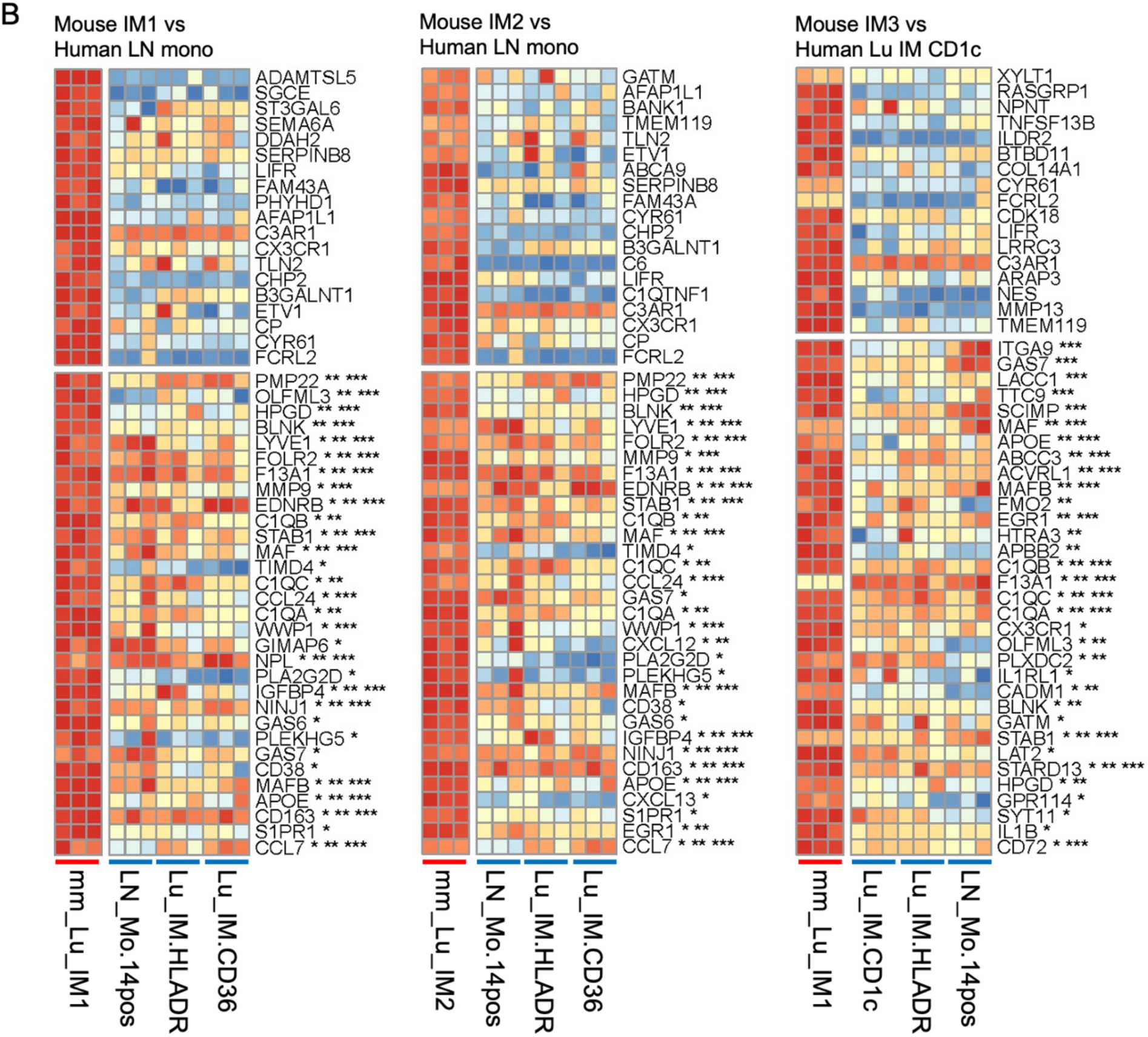
Corresponding human and mouse IM subtypes. Expression is shown for the first 50 ranked genes using the scaled data, as in Figure 5. Three replicates for the reference and three closest subtypes of the opposing species are shown. Data with all replicates can be found in Figure S10-S33 and is useful to consult for additional information about specific genes or cell types. Genes are annotated by whether they are also in the top 1000 marker genes in the three highest corresponding subtypes with *, ** or ***, respectively. A) human IMs. B) mouse IMs.

**Figure 7.**
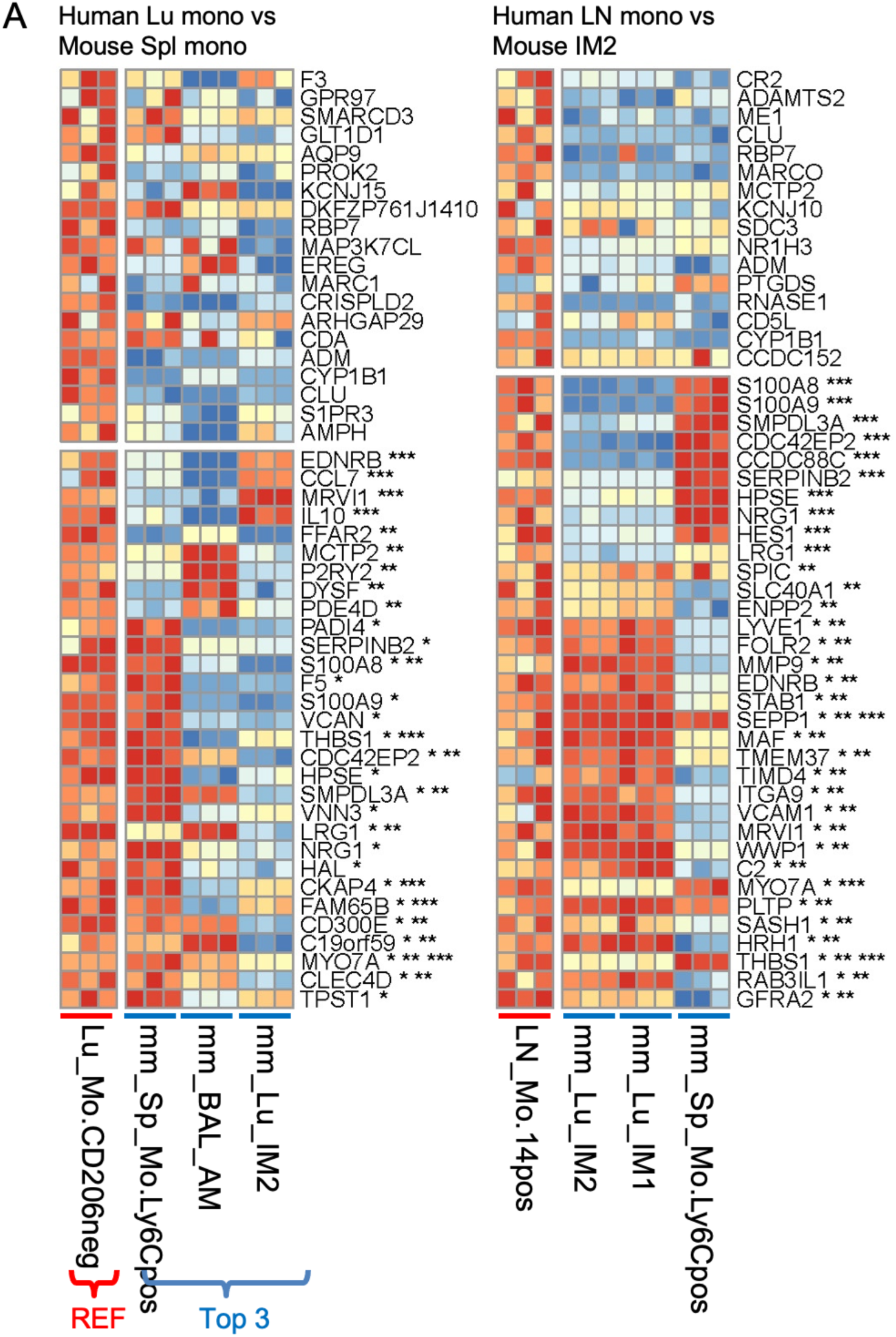
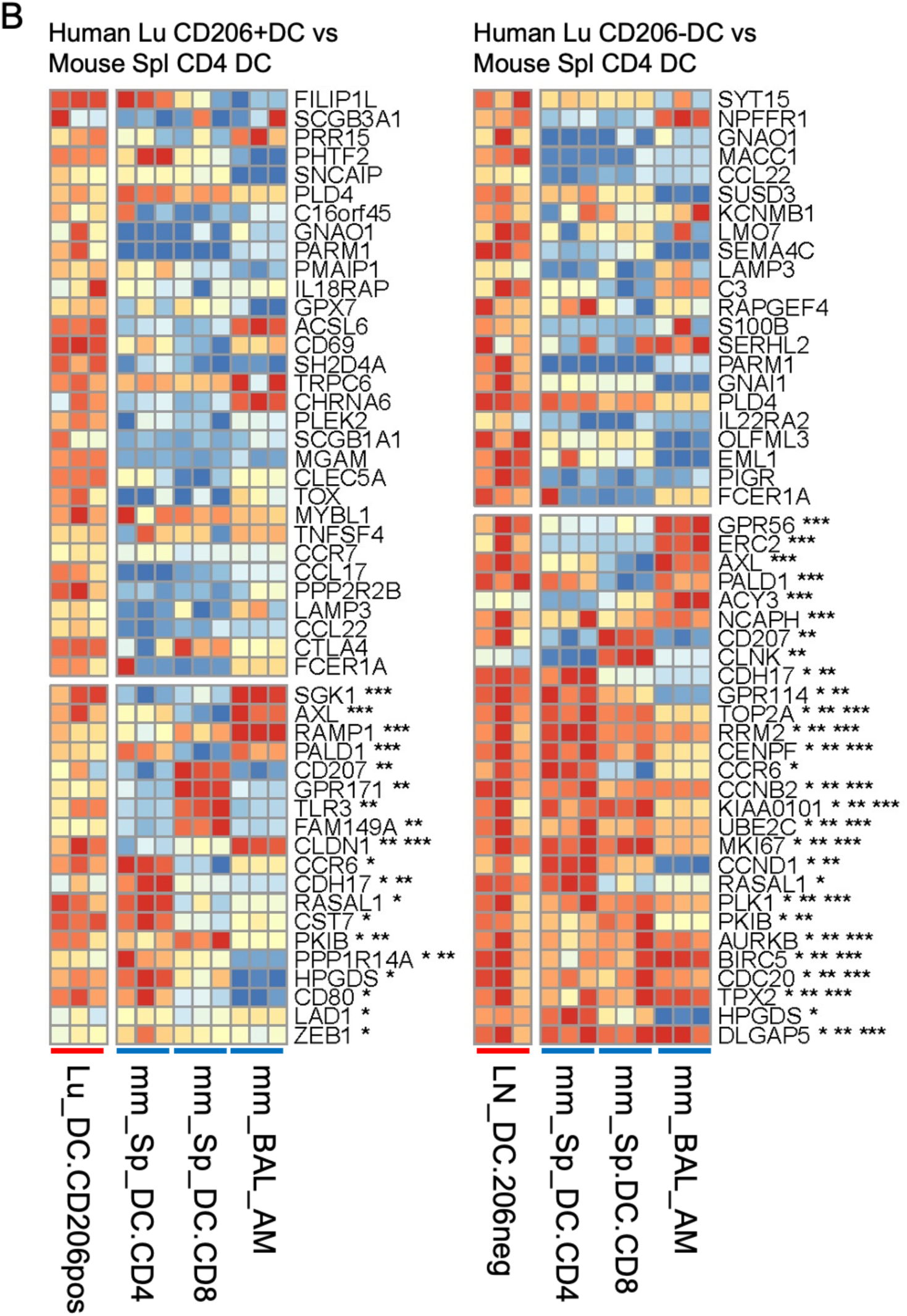
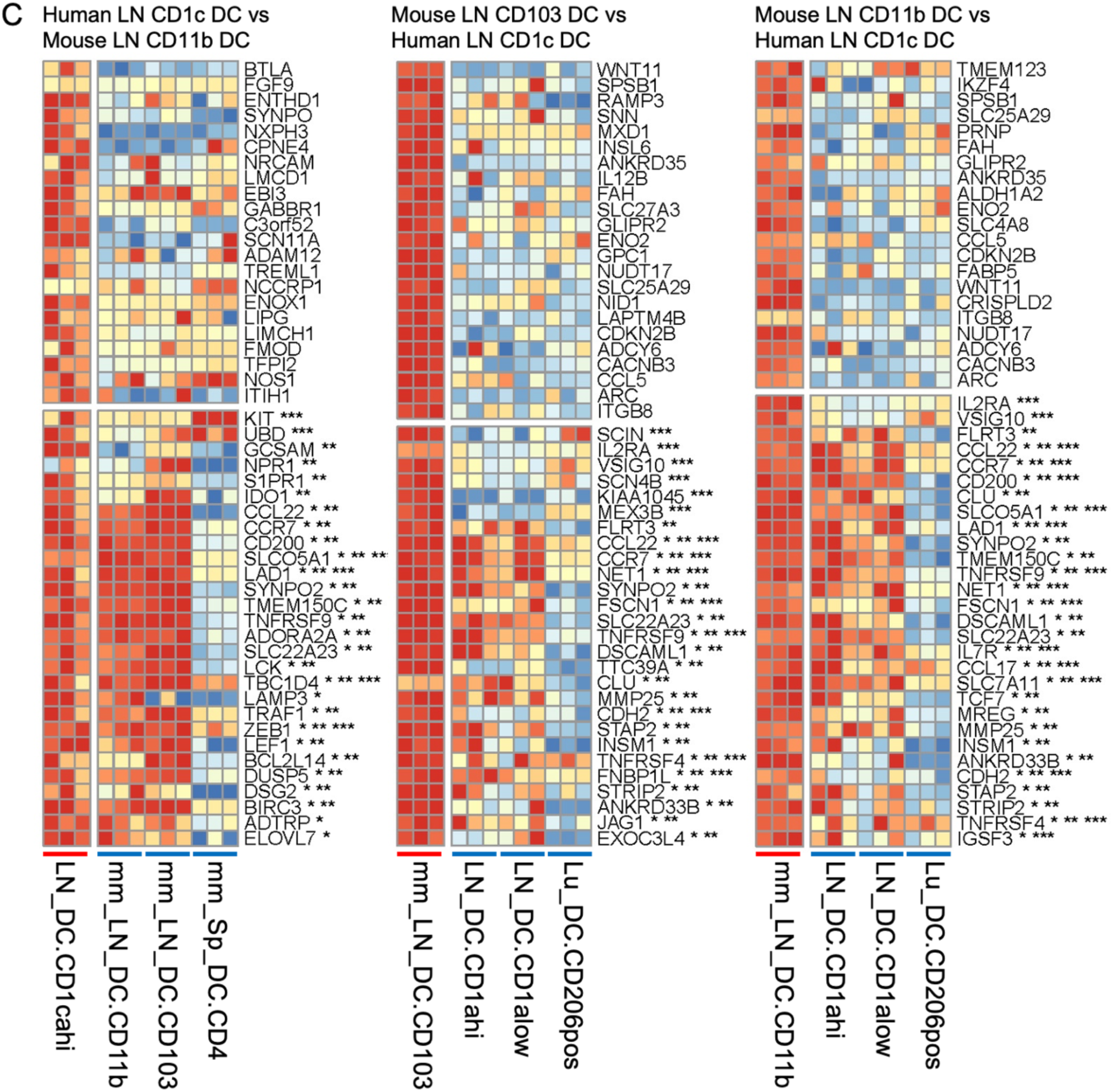
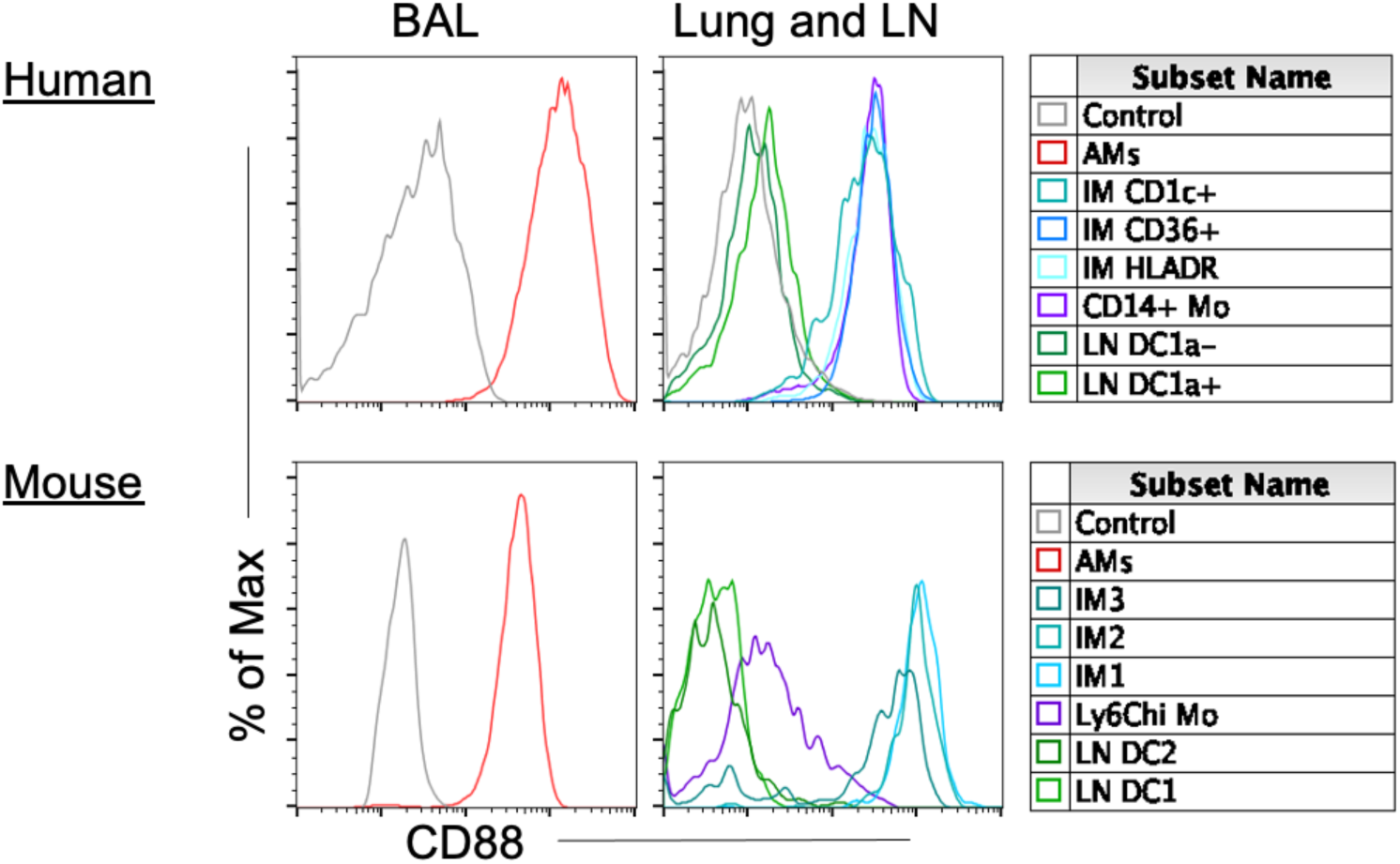
Corresponding human and mouse monocytes and DC subtypes; CD88 expression distinguishes monocyte/macrophages from DCs. Expression is shown for the first 50 ranked genes using the scaled data as in Figure 5. Three replicates for the reference and three closest subtypes of the opposing species are shown. Data with all replicates can be found in Figure S10-S33 and is useful to consult for additional information about specific genes or cell types. Genes are annotated by whether they are also in the top 1000 marker genes in the three highest corresponding subtypes with *, ** or ***, respectively. A) human monocytes B) human lung DCs C) human and mouse LN DCs. D) Histogram shows the protein expression of CD88 in human and mouse MPs in Figure 1.

As anticipated, signature genes that define these well-defined cell types are present such as *FABP4*, *MARCO and PPARG* for AMs (Group-1 AM, Figure 5B); *S100A8*, *S100A9,* and *CD14* for monocytes (Group-3 mono, Figure 5C); and *XCR1, CADM1,* and *CLEC9A*, along with a recently identified gene, *WDFY4*, required for cross-presentation for human blood Clec9a^+^ DCs and murine splenic CD8^+^ DCs (Group-6 DC1, Figure 5D) (Theisen et al., 2018). Although we outline genes known to be part of these human-murine homologs, it is important to note that many genes are not shared. For example, differences in gene expression for human-mouse AMs include *SERPING1, CES1, MME, IL17RB, and C1Q* (not shown), which are solely expressed in human AMs and not mouse AMs; whereas *SPP1*, *MMP8,* and *CARD11* are solely expressed in mouse AMs and not human AMs (Figure 5B). Human and mouse DC1, blood Clec9a^+^ and splenic CD8^+^ DCs, differentially express *IDO1* or *IDO*, *CD207* (langerin) and *CCR7* (Figure 5D).

### Conserved and distinct genes expressed in less well-defined human-mouse MPs

Since this method of analysis supports previously identified human-mouse MP homologs, we next assessed marker genes in less well-defined homologs: pulmonary IMs, LN monocytes, and pulmonary/LN DCs. Human IM CD36 and IM HLADR are best match with murine IM1, closely followed by murine IM2 (Group-2 IM, Figure 4B). The two murine IMs have a classical IM signature and share gene expression for *FOLR2, SEPP1, LYVE1, APOE, C1Q (not shown), CCL7,* and *CCL2* (Figure 6A). An example gene not shared is *SPP1,* highly expressed in human IMs (and murine AMs), but not murine IMs (Figure 6A).

Inversely, murine IM1 and IM2 match best with human LN monocytes followed by human IM HLADR and IM CD36 (Figure 6B). Unlike human blood and lung monocytes that correspond with murine splenic monocytes (Group-3 Mono), human LN monocytes reciprocally align best with murine IM1 and IM2 (Group-2 IM), followed by splenic monocytes (Figure 7A). Human LN monocytes share classical murine monocyte genes such as *S100A8*, *S100A9* and *CD14* (not shown), as well as mature IM genes such as *LYVE1*, *FOLR2*, *MMP9*, *F13A1*, *C1Q*, *MAFB, MAF, TIMD4,* and *STAB1* (Figure 7A).

Interestingly, murine IM3 matches best with human IM CD1c, where both MPs express inflammatory monocyte genes *IL1B, C1QB, C1QC,* and *F13A1* (Figure 6B, Group-5 DC2). However, human IM CD1c matches best with murine splenic CD4+ DCs (Figure 6A), based on their shared non-classical DC genes i.e., cytoskeletal genes, chemokine receptor 6, and cell cycle genes, *CD1K1* and *CDC20* (Figure 6A).

Similar to the alignment of monocytes and their implied state of maturation, human lung DCs align best with murine splenic CD4^+^ DCs compared to LN DCs (Figure 7B and Figure 3 and 7C, Group-5 DC2), expressing genes such as *CCR6*, *CD80*, *CD207*, and *AXL*. Consequently, human LN DCs align best with murine LN DCs (Figure 3 and 7C, Group-4 LN DCs) and express classical DC genes such as *ZEB1, CCL22, CCL17,* and *CCR7*, but not the human classical DC gene, *LAMP3* (Figure 7C). Overall, these in-depth correspondences allow one to investigate genes shared and divergent in both well-defined and less well-defined homologous pairs.

### A cell surface marker that allows a clear distinction between macrophage/monocytes and DCs in human and mouse

The rich expression compendium created in this study can be mined to identify new cell surface markers to isolate MP subtypes. In mice, the co-expression of F4/80 (EMR1) Ly6C, MerTK, and CD64 has helped distinguish DCs from macrophage and monocytes (Atif et al., 2018a, Yu et al., 2015, Misharin et al., 2013, Gautier et al., 2012). However, human monocytes do not have a Ly6C homolog, monocytes and macrophages express low levels of EMR1, and show variable expression of MerTK (perhaps due to the lack of a reliable antibody).

Analysis of our RNA-seq data identifies CD88 as a viable cell surface marker that can distinguish macrophages and monocytes from DCs in the tissue of humans and mice (Dutertre et al., 2019, Nakano et al., 2015). To validate this cross-species marker, histogram plots demonstrated that human and mouse AMs, IMs, and tissue monocytes express higher levels of CD88 compared to DCs (Figure 7D). All in all, CD88 is a commercially available cell surface marker that helps distinguishes MPs in humans and mice (Desch et al., 2015, Gautier et al., 2012).

## Discussion

The aim of this study was to investigate the homology between human and murine mononuclear phagocytes in order to aid in the translation of animal studies to human clinical uses. The dataset and analyses we provide are intended to improve the design of animal models mimicking human disease. Towards this goal, we created a gene expression compendium using RNA-seq analysis of MP subtypes from human and mouse, and a combination of computational methods to determine the closest cross-species counterparts. First, we identified marker gene signatures for each MP cell type and then compared these signatures against the MPs across species to identify homologous cell types. Overall, this dataset allowed us to explore the correspondence between human and murine MPs as well as to discover potential signature markers shared in common (or not) across human and murine MP subtypes.

One form of validation for our computational analysis, we analyzed cell types derived from our human blood and splenic MPs, and compared the populations from several recent single-cell RNA-seq (scRNA-seq) studies profiling human blood MPs (Villani et al., 2017, Dutertre et al., 2019), splenic MPs (Brown et al., 2019) or lung MPs (Travaglini et. al. bioRxiv https://doi.org/10.1101/742320) (Figure S9). Examining our expression data for the top 20 marker genes defined per cluster by the original publications, human blood Clec9a^+^ DCs consistently correspond to conventional DC1 populations and blood DC CD1c-cells correspond to plasmacytoid (pDC) populations. The monocyte subtypes express genes identified in monocyte clusters, while AMs and IMs express the top macrophage cluster genes. In general, our bulk-RNA-seq data of purified cell populations aligns favorably with the assumed cluster labels of scRNA-seq data, with the added advantage of robust measurement of thousands of genes across more MP cell types than previously published, and especially in the understudied LN MPs.

The validity and utility of our analysis for the identification of human-mouse MP counterparts was demonstrated by its *unbiased prediction of known homologs.* As expected, cross-species groupings include: 1) human and murine alveolar macrophages (Group-1); 2) human blood and murine splenic monocytes (Group-3); and 3) human blood Clec9a^+^ DCs and murine splenic CD8^+^ DCs (Group-6). These pairs aligned best with each other when compared with all other MPs across species (Ingersoll et al., 2010, Caminschi et al., 2008). These three pairs provided a useful reference for assessing the confidence with which our strategy could identify less well-defined MP subtype homologs. In addition, high degree of overlap between human blood and lung monocytes on one hand and mouse splenic monocytes on the other was reassuring, as it demonstrates a negligible effect on the transcriptome from cells extracted from blood versus digested tissue.

We also observed a preserved general distinction between MP lineages in both human and mouse. For example, Groups 1, 2, and 3 contain exclusively macrophages (AMs and IMs) and monocytes and whilst groups 4, 5, and 6 contain DCs (Figure 3, Figure 4B), with two exceptions. The first exception is human LN monocytes in the IM-enriched Group-2, which as previously stated represents a more matured monocyte expressing both classical monocyte and differentiated macrophage genes. The second exception is human lung IM CD1c^+^, which aligns more closely with murine splenic CD4^+^ DCs than its expected counterpart mouse lung IM3; although, conversely, mouse IM3 aligns best with lung IM CD1c^+^ (Figure 4B).

Interestingly, human blood DCs and mouse splenic DCs clustered away from human and mouse LN DCs, perhaps due to tissue imprinting. One distinction we noticed is the expression of CCR7, which is a hallmark feature of DCs. Human blood DCs lack CCR7, a receptor required for lymphatic migration and localization in the T cell zone of LNs (Randolph et al., 2005). This is possibly because the cells are immature and require additional endothelial-tissue interaction to up-regulate CCR7. Another observation is that human DC subtypes differentially express CD62L, a transmigration molecule used by circulating monocytes and preDCs to enter tissue and LNs (Liu et al., 2009, Jakubzick et al., 2013, Leon and Ardavin, 2008). CD62L is highly expressed in human blood DCs, but not in tissue DCs, suggesting that once in tissue this gene is no longer required. Thus, one possibility could be that human blood DCs may give rise to human tissue DCs. An alternative possibility is that human blood DCs and tissue DCs share the same precursor cell, which gives rise to DCs in different compartments. There is a precedent for this in mice: Murine lymphoid resident DCs do not give rise to migratory tissue DCs but share the same precursor cell, where the former DCs precusor entered the LN via the HEV and the latter DC migrated from the tissue into its draining LN via afferent lymphatics.

One surprising finding was the relatively small effect of DC lineage on gene expression patterns. In mice, there is a strong distinction between DC1 (Batf3-dependent) and DC2 (Irf4-dependent) lineages, as seen for CD8^+^ (DC1) versus CD4^+^ (DC2) splenic DCs. The existence of two DC lineages in humans is supported by the alignment of blood Clec9a^+^ DC with murine CD8^+^ splenic DC (Group 6), compared with CD4^+^ splenic DCs. However, the separation of human lung or LN DCs based on DC lineage appears to be masked by the stronger effects of tissue imprinting, as evidence by the distinct clustering of the LN DC (Group 4) from lung DCs (Group 5) and human blood/splenic DC1 (Group 6). In addition, we originally separated LN DCs based on CD1a expression because we consistently observed CD1a^hi^ and CD1a^lo^ CD1c^+^ lung-draining LN DCs in over 50 individuals, making the assumption that they might be distinct cell types. However, it is now clear that although the sorted human LN DCs are distinct at the CD1a protein level, these populations are not transcriptionally distinct. Further investigations on how to subset DEC205^+^CD1c^+^ lung-draining LN DCs using different cell surface markers is needed. More importantly, human LN DCs are understudied. Even though we were unable to decipher more than one LN DC cluster, there is a wealth of genetic information provided in this dataset for future follow up. For instance, CCL17 and CCL22 expression is restricted to DC populations in lung and LNs, and is not expressed in human blood DCs (Alferink et al., 2003, Stutte et al., 2010, Rapp et al., 2019, Vulcano et al., 2001). This is also true in mice, where these chemokines distinguish DCs from other immune cell types and are important for T cell recruitment and interactions (Rapp et al., 2019).

Overall, the overlap among the top 1000 genes for corresponding human and mouse MPs of the same type was low (13-28%, Figure S6B). This does point to the need to identify species-specific marker genes, and not assume that a gene identified in one species will be an adequate marker in another species. This applies to both translational (mouse to human) and reverse translational (human to mouse) studies. However, it is important to note that this does not mean that the species have completely different transcriptional profiles overall. To qualify as a marker gene for a given MP subtype, a gene had to be expressed primarily in just that one single subtype. Genes expressed in common across multiple closely related subtypes were not included in the overlap analyses (since these genes are not unique to one subtype). An alternative approach would be to detect the broader differences between monocytes/macrophages versus DC lineages overall, including all genes. Broader inclusion would likely yield a much higher percentage overlap across all MP subtypes, and estimating this overlap may be useful for some applications. However, it would require a different analysis and is beyond the scope of the present study, as our goal was to focus on the subset of genes likely to be useful as markers for particular MP subtypes.

Finally, in our preliminary investigation, we identified CD88 expression to be highly predictive of macrophage/monocytes compared to DCs in both human and mouse. Fortunately, there are commercially available antibodies against CD88 (and CD64, shown previously) that work well for separation of monocytes/macrophages from DCs in human and mouse (Desch et al., 2015, Nakano et al., 2015, Gautier et al., 2012, Jakubzick et al., 2013). In conclusion, this study provides a platform to aid in the design of future translational studies at the MP subtype level and at the gene level. Awareness of cross-species homology, or lack thereof, can save a great amount of time and expense in medical research.

## Acknowledgments

This manuscript is dedicated to the lung donors and their families that enabled this research. Thanks to Donna Bratton, Peter Henson and Elizabeth Castel for project discussions and technical support isolating human myeloid cells. Funding: NIH R01 HL135001 and R01 HL115334.

## Authors contributions

SML, SLG, and CVJ prepared the manuscript and designed the experiments. SLG, AT and SMA executed sorting and flow cytometry. SML, TD and BV executed the bioinformatics and biostatistics data. All authors provided intellectual input and critical feedback.

## STAR Methods

### RESOURCE AVAILABILITY

#### Lead contact

Further information and requests of resources and reagents should be directed to and will be fulfilled by Dr. Claudia Jakubzick.

#### Materials Availability

This study did not generate new unique reagents.

#### Data and Code availability

Transcriptome data from murine mononuclear phagocytes can be accessed from the NCBI Gene Expression Omnibus via accession numbers GSE132911 (monocyte and DCs) and GSE94135 (AMs and IMs). To protect donor anonymity human RNAseq data have not been deposited in a public repository. Requests for access to human sequencing data may be made to the lead contact.

### EXPERIMENTAL MODEL AND SUBJECT DETAILS

#### Human subjects

We received de-identified human lungs that were not used for organ-transplantation from the International Institute for the Advancement of Medicine (Edison, NJ, USA) and the University of Colorado Donor Alliance. We selected donors without a history of chronic lung disease and with reasonable lung function with a PaO2/FiO2 ratio of >225, a clinical history and X-ray that did not indicate infection, and limited time on a ventilator. Lungs from 7 individuals were processed for RNA sequencing; the mean donor age was 35.8 years (SD 24.7 years), 3 were female and 4 were male. 4/7 donors were non-smokers (defined as never having smoked) 2 were light smokers (less than a pack a year) and 1 was a former smoker. Additionally, blood was drawn from 3 healthy, non-smoking volunteers, all female with a mean age of 30 years (SD 4.6). Additional demographic information is shown in Table S2. 10 different mononuclear phagocyte (MP) populations were sorted from each lung donor, and 5 different populations were sorted from each blood donor, as outlined in Figure 1. Note, not every donor provided all 15 MP cell types. After data quality control analysis, this resulted in 73 total human MP samples. The Committee for the Protection of Human Subjects at National Jewish Health approved this research.

#### Animals

Naive female C57BL/6 mice were obtained from Charles River Laboratories. Mice were group housed in a specific pathogen-free environment at National Jewish Health (Denver, CO), and used at 6–10 weeks of age, in accordance with protocols approved by the Institutional Animal Care and Use Committee. Each sample represents pooled cells from 5 mice; different cohorts of mice were used to isolate lungs, LLNs and spleens.

### METHOD DETAILS

#### BAL, lung, lymph node, and PBMC cell preparation

Lungs were removed en bloc in the operating room and included the trachea, LNs and pulmonary vessels. Pulmonary arteries were perfused in the operating room with cold histidine-tryptophan-ketoglutarate (HTK) solution to preserve endothelial cell function and prevent intravascular clot formation. The lungs were submerged in HTK, and immediately shipped on ice. All lungs were processed within 24 hours of removal. The lungs were visually inspected for lesions or masses and were eliminated from the study if grossly abnormal. Peribronchial LNs were identified and removed. Procedure additionally described in (Desch et al., 2015, Gibbings and Jakubzick, 2018). *Bronchoalveolar lavage* (*BAL*) was performed on the right middle lobe or lingula by completely filling the lobe three times with PBS and 5mM EDTA and then 3 times with PBS alone. After each instillation, lavage fluid was drained from the lung, collected and pooled.

*Lung* tissue digestion was performed by inflating the lobe with 4U/mL elastase solution (Worthington Biochem) in PBS containing Ca2+, Mg2+, HEPES and dextrose. Airways were clamped to prevent leakage of enzyme and the tissue was incubated at 37°C for 40 minutes. Manual disruption of the tissue was performed, firstly with scissors and secondly with a food processor in buffer containing fetal bovine serum. The resulting tissue homogenate was filtered through a 100μ nylon filter membrane to create a single-cell suspension. Density gradient centrifugation using 30% and 65% Percoll (Sigma Aldrich) was used to deplete red blood cells and dead cells.

*Lymph nodes* were identified and dissected from lung tissue, pooled, minced and enzymatically digested with 2.5 mg/mL collagenase D (Roche/Sigma) for 30 minutes at 37°C. Digested tissue was collected and filtered through a 100μ nylon filter membrane to create a single cell suspension.

*Peripheral blood mononuclear cell (PBMC) isolation,* blood was obtained by venipuncture from healthy adults as per IRB approved protocol. CPT tubes (BD) were used to separate PBMC from other blood components according to manufacturer’s instructions.

#### Murine Lung, LLN and spleen MP isolation

Mice were euthanized by CO_2_ asphyxiation and lungs were perfused with cold PBS via the heart. Lungs were removed, minced finely with scissors and subjected to enzymatic digestion with 0.5mg/mL Liberase (Roche) for 30 minutes at 37°C as reported previously (Gibbings et al., 2017). Mediastinal lymph nodes or spleens were removed, minced, and digested by incubation with 2.5mg/mL Collagenase D for 30 minutes at 37°C. Single cell suspensions were prepared by manual homogenization and filtration through a 100μ nylon filter membrane. Mouse lung macrophages were labeled with anti-mouse MerTK and enriched with anti-biotin microbeads as previously reported. Mouse lymph node and spleen DCs were enriched with anti-mouse CD11c microbeads and spleen monocytes were enriched using anti-mouse CD11b microbeads according to the manufacturer’s protocol (Miltenyi Biotec).

#### Mononuclear phagocyte enrichment

Cells of interest were positively enriched prior to FACS sorting using Miltenyi microbeads according to manufacturer’s protocols. Human blood monocytes and dendritic cells were enriched from PBMCs using biotin-conjugated anti-CD141 and anti-CD1c in combination with anti-biotin microbeads. Enrichment of human lung and lymph node cells was achieved using PE conjugated anti-human CD1c and biotin-conjugated anti-human CD11c antibodies in combination with anti-PE and anti-biotin microbeads.

#### FACS sorting and flow cytometry

Human BAL, enriched blood, lung and lymph node cells were re-suspended in PBS with 2mM EDTA, 0.1% BSA and 10% pooled human serum, and stained with fluorescently conjugated antibodies as described in the key resources table. Dead cells were excluded by uptake of 4′,6-diamidino-2-phenylindole (DAPI, Invitrogen/ Thermo Fisher Scientific). Cells were either sorted using a FACS Aria Fusion (BD) as outlined in Figure 1 for RNAseq analysis or analyzed using an LSRII or LSR Fortessa (BD).

#### Human RNA-seq data

Isolated mononuclear phagocytes were immediately re-suspended in RLT buffer for RNA extraction using Qiashredder columns and the RNeasy micro kit (Qiagen) per manufacturer’s instructions. Isolated total RNA was processed for next-generation sequencing (NGS) library construction as developed in the NJH Genomics Facility for analysis with a HiSeq 2500 (Illumina San Diego, CA, USA). A Clontech SMART-Seq® v4 Ultra® Low Input RNA Kit for Sequencing (Mountain View, CA, USA) and Nextera XT (Illumina San Diego, CA, USA) kit were used. Briefly, library construction started from isolation of total RNA species, followed by SMARTer 1st strand cDNA synthesis, full length dscDNA amplification by LD-PCR, followed by purification and validation. After that, the samples were taken to the Nextera XT protocol where the sample is simultaneously fragmented and tagged with adapters followed by a limited cycle PCR that adds indexes. Once validated, the barcoded-pooled libraries were sequenced using 1×50bp chemistry on the HiSeq 2500 as routinely performed by the NJH Genomics Facility.

#### RNA-Seq data generation

For human samples, FASTQ files were generated with the Illumina bcl2fastq converter (version 2.17). Nextera TruSight adapters were trimmed, and degenerate bases at the 3’- end, as well as highly degenerate reads were removed using skewer (version 0.2.2), which by default removes reads with a remaining length of less than 18 nt. For the mouse samples, basecalling, barcode demultiplexing and adapter trimming was performed using the Ion Torrent Suite software, and reads less than 30 nt in length were discarded. The quality of the reads was assessed using FastQC (version 0.11.5). The preprocessed sequence reads were mapped to the genome of the respective species (canonical chromosomes of release hg 19 for human and mm10 for mouse, respectively) using STAR (Dobin et al., 2013). Reads mapping uniquely to each gene of the Ensembl 75 annotation of the respective species were quantified with the featureCounts program from the subread software package (version 1.5.2, using parameters -s 0 -O --fracOverlap 0.5 (Liao et al., 2014).

### QUANTIFICATION AND STATISTICAL ANALYSIS

#### Cross-species RNA-seq data

A cross-species data set of RNA-seq reads was obtained for all autosomal genes with a 1:1 homology between human and mouse, as determined Ensembl BioMart version 75 (Homo sapiens genes, Homologues subsection, Mouse Orthologs). Sex chromosome genes were omitted to avoid a strong effect of donor gender. The resulting cross-species data consisted of 11,959 genes across 93 samples total. For visualization purposes, gene expression levels that had subject effects regressed out were obtained for both the human and mouse samples, as follows. The original counts were transformed to a continuous measure using the variance stabilizing transformation (VST) available from the DESeq2 R package version V1.20.0.(Love et al., 2014). Using these transformed values, linear mixed models were fit to each gene with the nlme R package version V3.1-137. These models included an intercept only for fixed effects along with a random intercept for each subject. The expression values with subject regressed out were obtained by adding the overall intercept with the residual error terms (thus excluding the random intercepts). For the small number of genes where the linear mixed models had a singular fit (i.e. the random intercept variance was estimated to be 0), the original expression values were retained. This was performed separately for the human and mouse samples. To further account for differences in expression levels across species, the VST subject-regressed data was quantile-normalized using the preprocessCore package (version 1.40.0) in R version 3.4.2 (Figure S3). The quantile-normalized data was used in all subsequent analyses, except the Z-score analysis described below.

#### Alignment and visualization of human-murine MP data

Human and murine quantile-normalized gene-expression data were aligned and jointly visualized using the Seurat pipeline, originally developed for single-cell sequencing and previously demonstrated to integrate human and murine single-cell data. The standard pipeline was applied here to the human and murine MP data using the Seurat 2.3.2 package for R version 3.4.2 (Satija et al., 2015). Briefly, the data were first normalized using the default methods in Seurat to yield natural-log transformed data. Then a set of 885 highly variable genes (hvg) was identified as shared among the 2000 most highly variable human genes and 2000 most highly variable mouse genes. The data were then scaled using the standard Z-score scaling before further dimensionality reduction. Canonical Correlation Analysis (CCA) was applied to the scaled hvg set to identify a lower dimensional space of CC vectors that maximized the shared correlational structure across human and mouse data sets. Using the MetageneBicorPlot function to display the correlation strength for each CC vector, the optimal number of 12 CC vectors was identified for use in the dynamic time warping alignment step. The resulting single, aligned, low-dimensional space representing the integrated human and mouse data was visualized in 2-D using t-Distributed Stochastic Neighbor Embedding (t-SNE). (Figures 3, and S4).

#### Identifying cell-type specific marker genes

Marker genes were identified as those significantly over-expressed primarily in one MP subtype versus all others within a species. Over-expression was quantified using the quantile normalized VST counts. The median expression was computed for each gene in each subtype within a species and then scaled by the median of those expression values across all samples within the species. Scaling by the median expression helps distinguish between genes highly expressed primarily in a minority of subtypes from genes like housekeeping genes that are highly expressed in all subtypes.

Significance of over-expression in one MP subtype was determined by calculating a Z-score for each gene for each subtype of interest as follows. RNA-Seq counts from genes that had a human-mouse equivalent were filtered to include only genes with at least 3 samples having more than one read count per million aligned reads. Next, the counts were transformed to continuous values using the VST from the DESeq2 R package. For each gene, a linear mixed model using subtype for the fixed effects and a random intercept for subjects was fit using the lmerTest R package version 3.1-0 (Kuznetsova et al., 2017). An overall F-test, using Satterthwaite’s degrees of freedom method, to determine which genes showed any evidence for differences in expression between the subtypes was performed, and then these p-values were adjusted using the Benjamini-Hochberg method. For genes with adjusted p-values ≤ 0.05 for the overall F-test, a meta-analysis approach was used to evaluate potential signature genes for each different MP subtype. For the subtype of interest, all pairwise comparisons were done using linear contrasts and t-tests (again with Satterthwaite’s degrees of freedom) using the linear mixed model described above. The raw p-values were then combined using the Z-score method where pi was used if the effect size for that pairwise comparison was positive (i.e. the expression was higher in the subtype of interest) or max (pi, 1-pi) was used if the expression was lower in the subtype of interest. The same process was repeated for the mouse samples. A higher positive Z-score indicates that the gene is expressed higher in the subtype of interest than the other subtypes, on average.

A final marker score for each candidate gene per subtype was computed as the product of the median scaled expression data and the Z-score using that subtype as the subtype of interest. In this way, genes that were both highly and preferentially expressed in a single subtype would be ranked more highly by the marker score. Note, this strategy discounts genes like housekeeping genes that may be highly expressed in all subtypes but not preferentially in a given subtype, as well as genes that could be marker genes for multiple cell types.

#### Identifying human-mouse MP analogs

A popular method for determining the corresponding MP subtype between species is to minimize distances or maximize correlations between sample groups using the quantile-normalized gene expression vectors, or in the lower dimensional projection defined by the aligned CC vectors. However, with so few replicates per sample group and a difference in gene variance structure across species, both these methods were found to be very sensitive to outliers and data scaling (data not shown). Moreover, determining the corresponding cell types by maximizing correlation between samples is highly dependent on the set of genes over which the correlation is computed (Figure S5). Whether using all genes (Figure S5A) or only the highly variable genes (hvg) to avoid flat or non-expressed genes dominating the correlation (Figure S5B), all samples are nearly equally correlated, with little discrimination. Moreover, the pairings which maximize correlation using all genes do not always agree with the pairings using the hvg. Another alternative to determining homologous subtypes is to intersect marker gene lists and report as the best match that which has the largest marker gene overlap (Figure S6). However, for the marker gene determination used here (positive Z-score and p.adj>0.05), ∼4,000-7,000 of the initial 11,959 genes were chosen as potential marker genes per cell type. Given the large marker gene set size relative to the total set of genes, comparing any two sets would necessary overlap of nearly 50% (Figure S6A). Restricting the overlap analysis to fixed sized sets of the top ranked marker genes per candidate pair results in different ‘best’ matches depending on the size of the marker set (Figure S6B). To compute an unbiased correspondence between human and murine MP, the data were analyzed instead based on a summary value of average overlap percentage when considering ordered marker set sizes from 1 to 1000 identified within each species. The set of marker genes for a reference MP subtype in one species was ranked in decreasing order of marker score and compared to the similarly ranked marker genes in the subtypes of the other species. The proportion of genes in common between the reference marker ranked list and each candidate cell type ranked list was plotted against the list size as a Correspondence-at-the-Top (CAT) plot. (Figures 4A, S7 and S8). Due to the ranking criterion, higher agreement at smaller list sizes, i.e., the top of the list, indicates two subtypes agree strongly on genes most highly and preferentially expressed in a single subtype per species, i.e. marker genes unique to the cell type. For each candidate subtype cross-species pair, the overlap percentage was averaged across all list sizes from 1 to 1000 (Figure 4B). A high mean CAT percent agreement indicates that the pair consistently agrees on strong overlap, regardless of the marker set size (from 1 to 1000). The cross-species analog was chosen as that which maximized the mean CAT percentage agreement across all list sizes from 1 to 1000 (Figure 4B).

#### Comparison to previous DC and monocyte signature gene sets

The signature gene sets defined by Villani et al., (Villani et al., 2017) for the six DC subsets and for four monocyte subsets. Genes not appearing in our dataset were first removed and the top 20 genes per subset were identified using the reported AUC value to rank the genes (AUC column, Tables 1, S2 and S5 in (Villani et al., 2017) paper). Signature gene sets defined by Dutertre et al., (Dutertre et al., 2019) appearing in our dataset or occurring as multiples in their signatures were first removed and then the top 20 genes per subset were defined based on decreasing order of average log fold change (avg_logFC column, Table S2 in (Dutertre et al., 2019) paper). The correspondence of cluster number to cluster identity was established using the signature genes highlighted for each cluster in Dutertre et al. Signature gene sets from Brown et al., (Brown et al., 2019); were created from the normalized count data available the Gene Expression Omnibus (GEO) accession GSE137710. Following the methods described in Brown et al., a score for each gene for each subset was computed as the product of the earth mover’s distance (EMD) and the area under the ROC curve (AUC) using the gene to distinguish between cells within the cluster versus all cells outside the cluster. Genes not appearing in our dataset were removed and the top 20 genes per DC subset were defined based on decreasing order of the gene score. The top scoring genes were verified as the genes annotated in Brown et al. MP signature gene sets were defined by Travaglini et al., (https://www.biorxiv.org/content/10.1101/742320v2). Genes not appearing in our dataset were first removed and the top 20 genes per subset were identified based on decreasing order of the average log fold change (avg_logFC column, Table S4 in Travaglini et al., paper).

### KEY RESOURCES TABLE

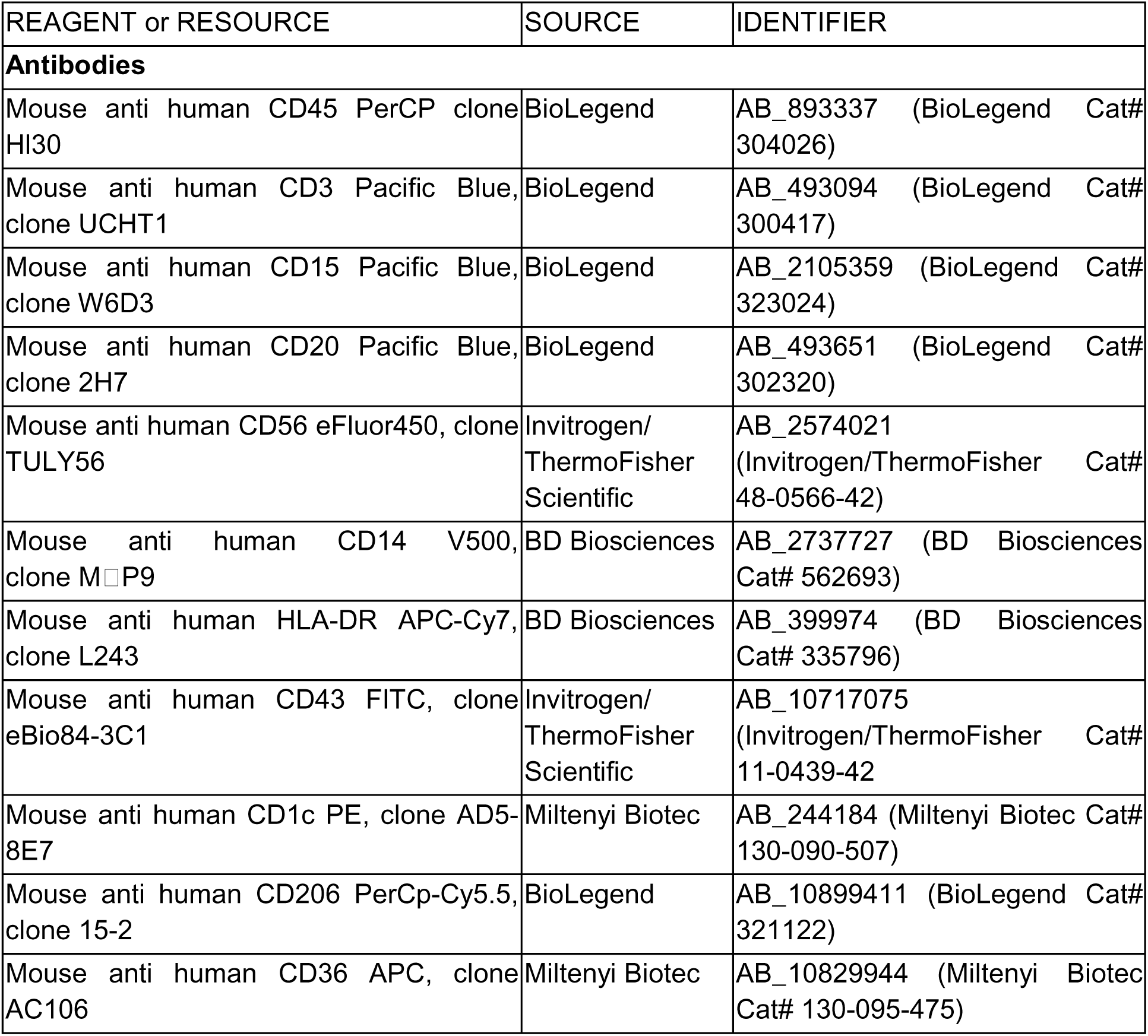
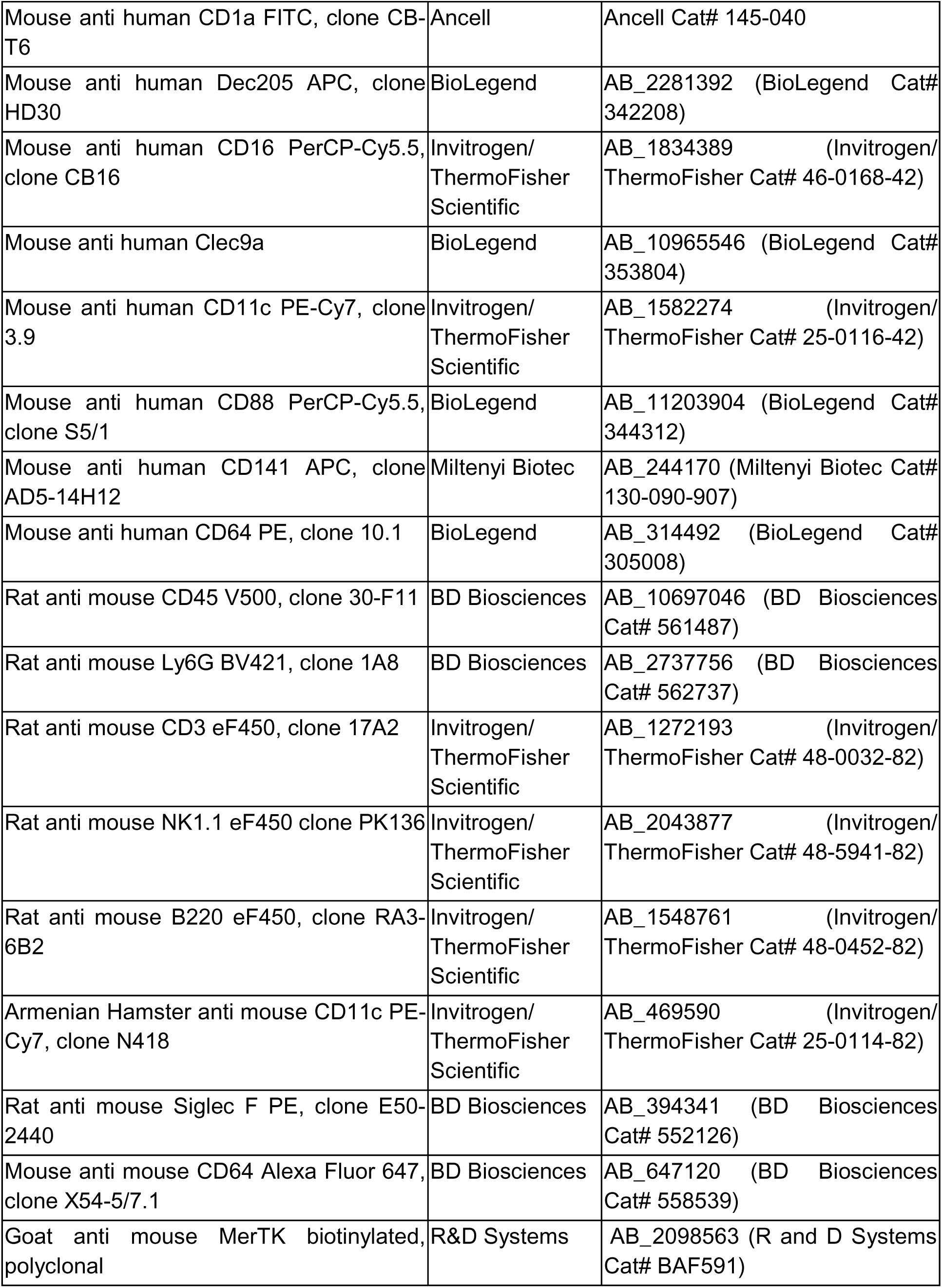
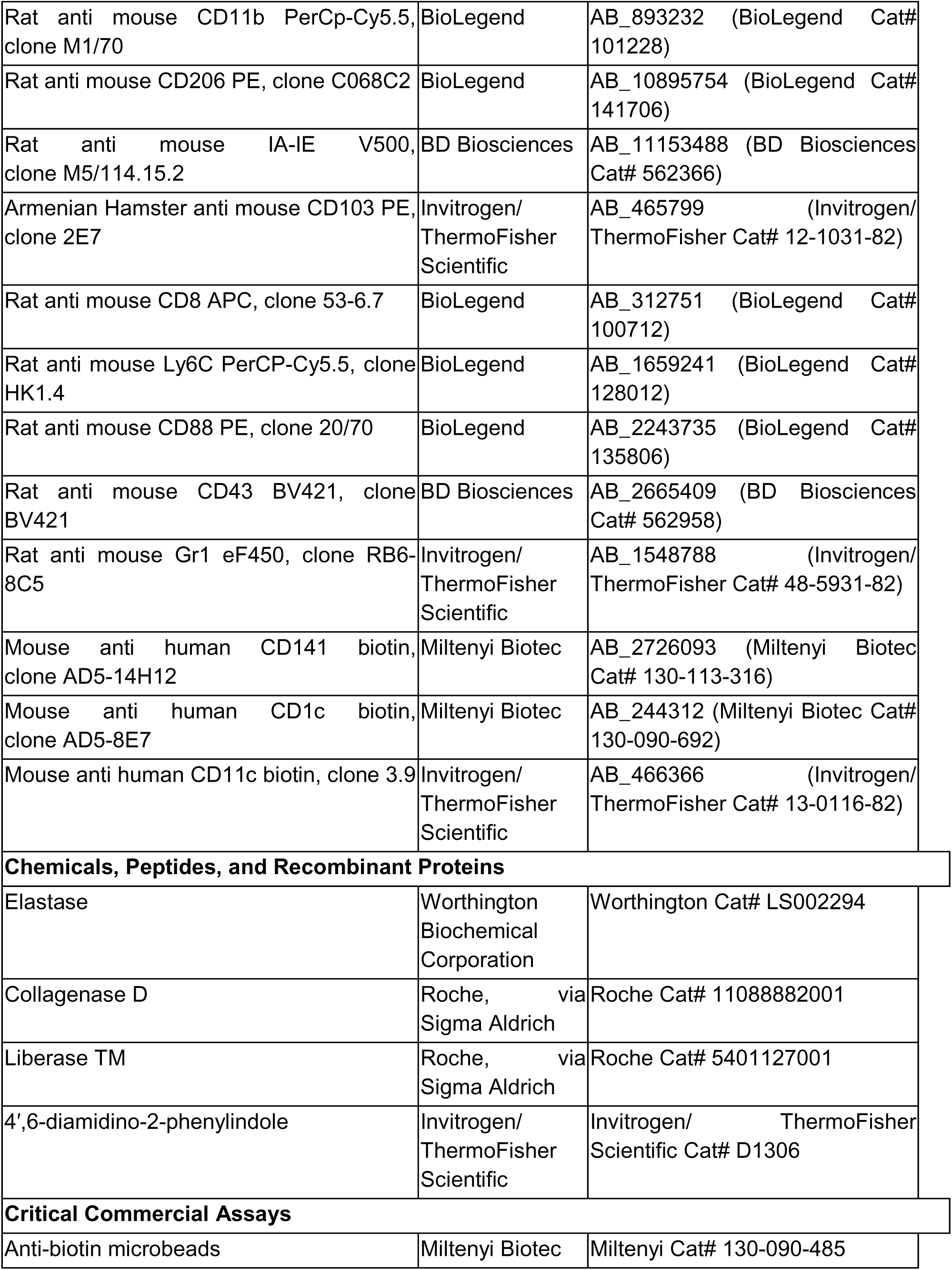
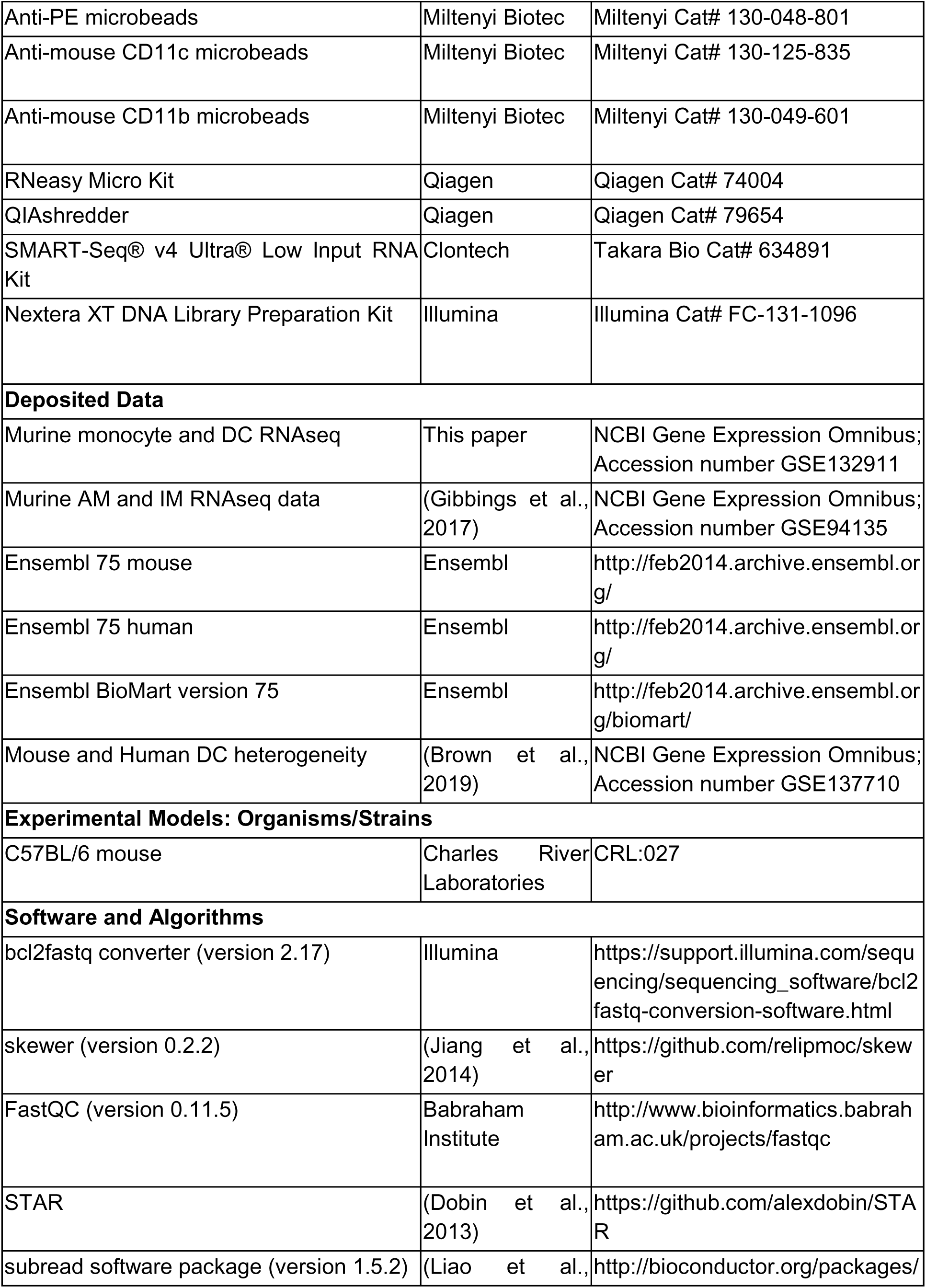
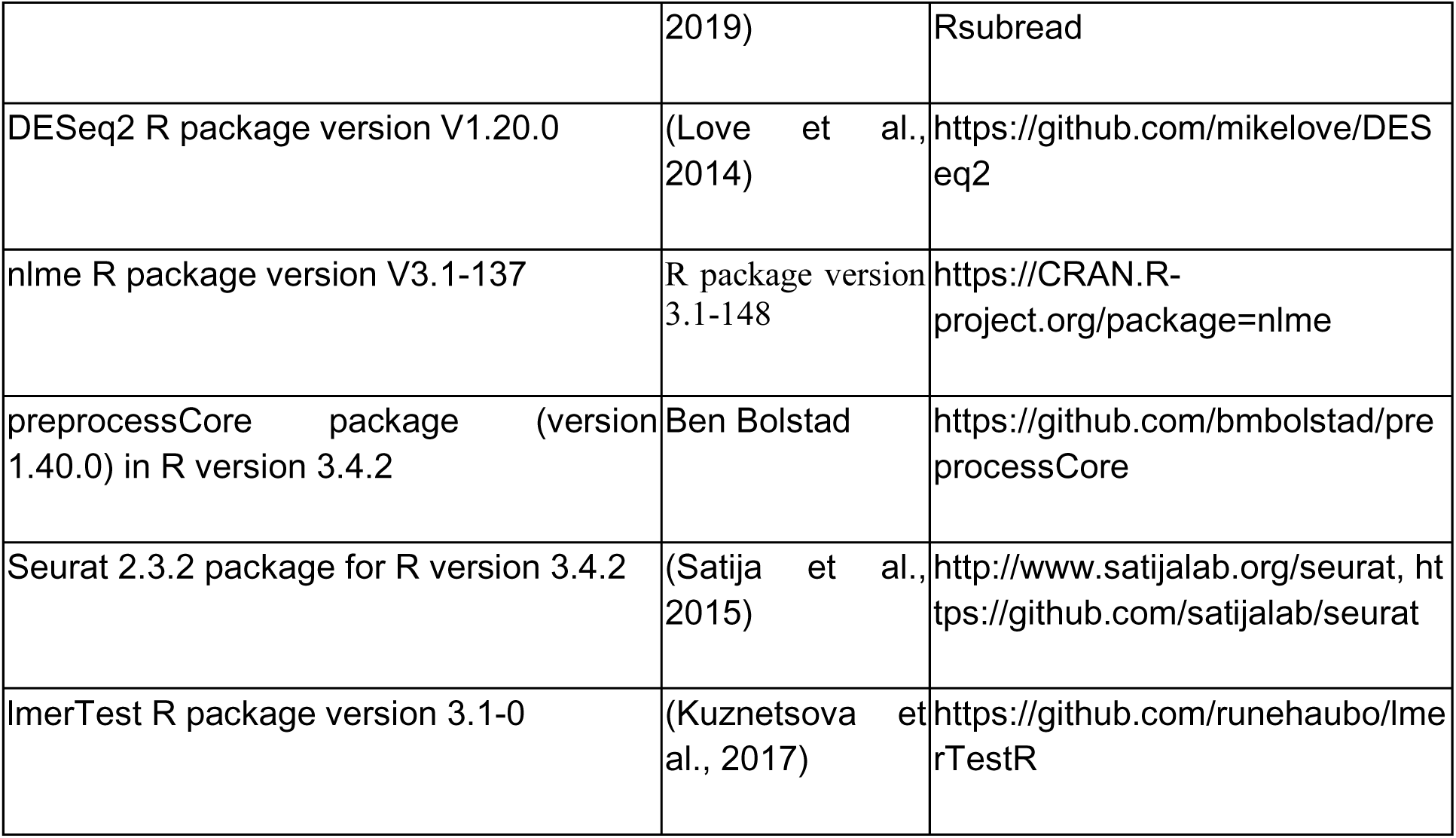

## Supplemental Information

**Table S1.**
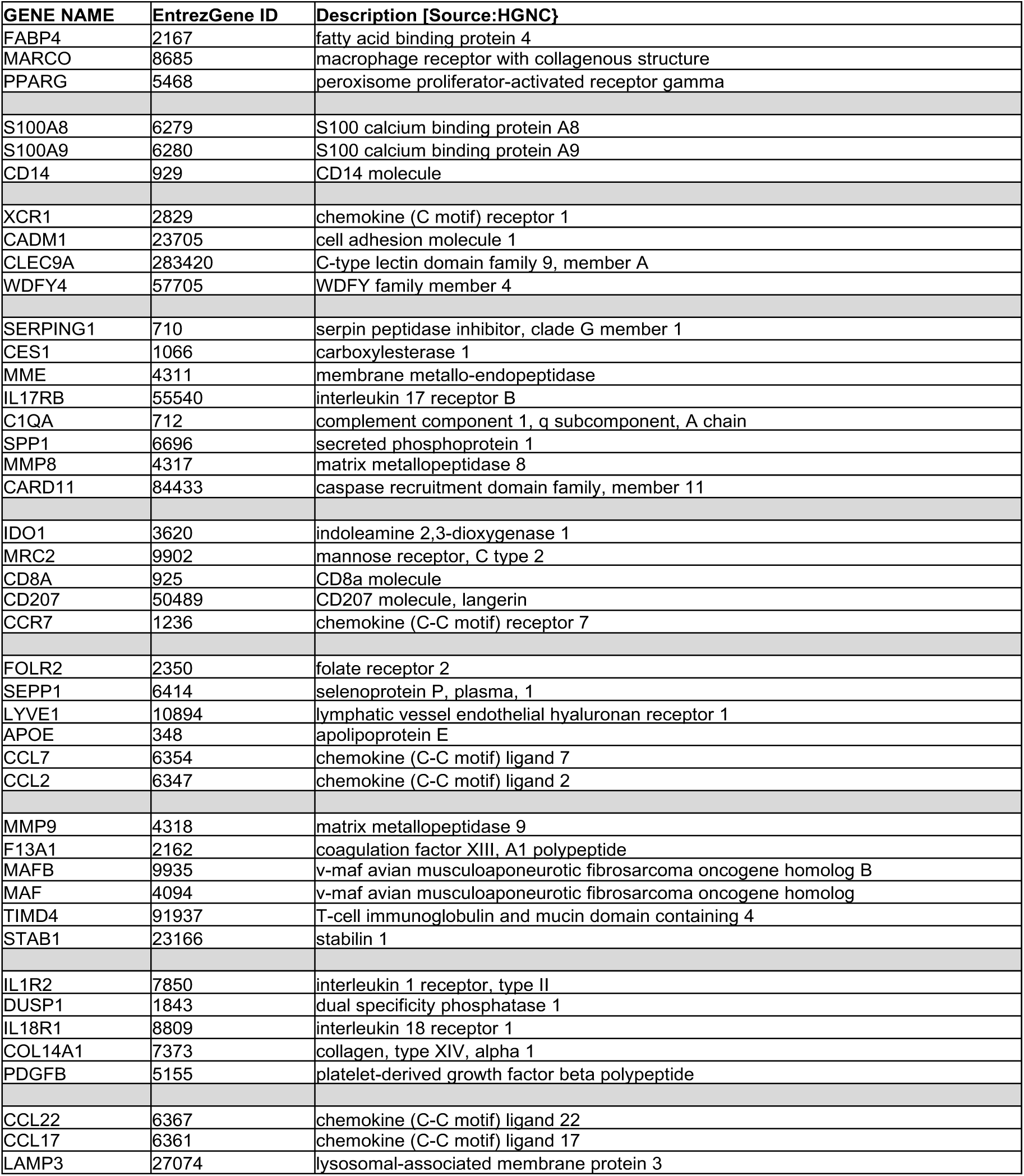
EntrezGene ID and description of MP marker genes

**Table S2.**
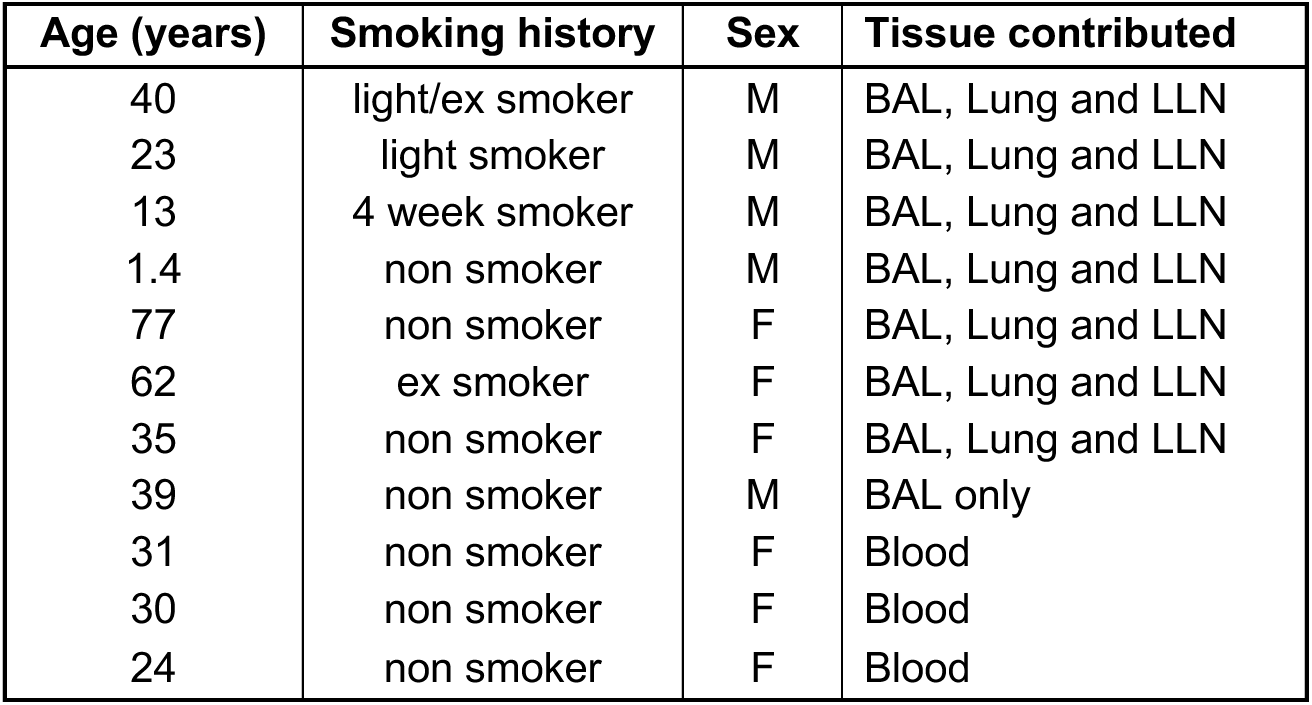
Demographics for RNA-seq

**Figure S1.**
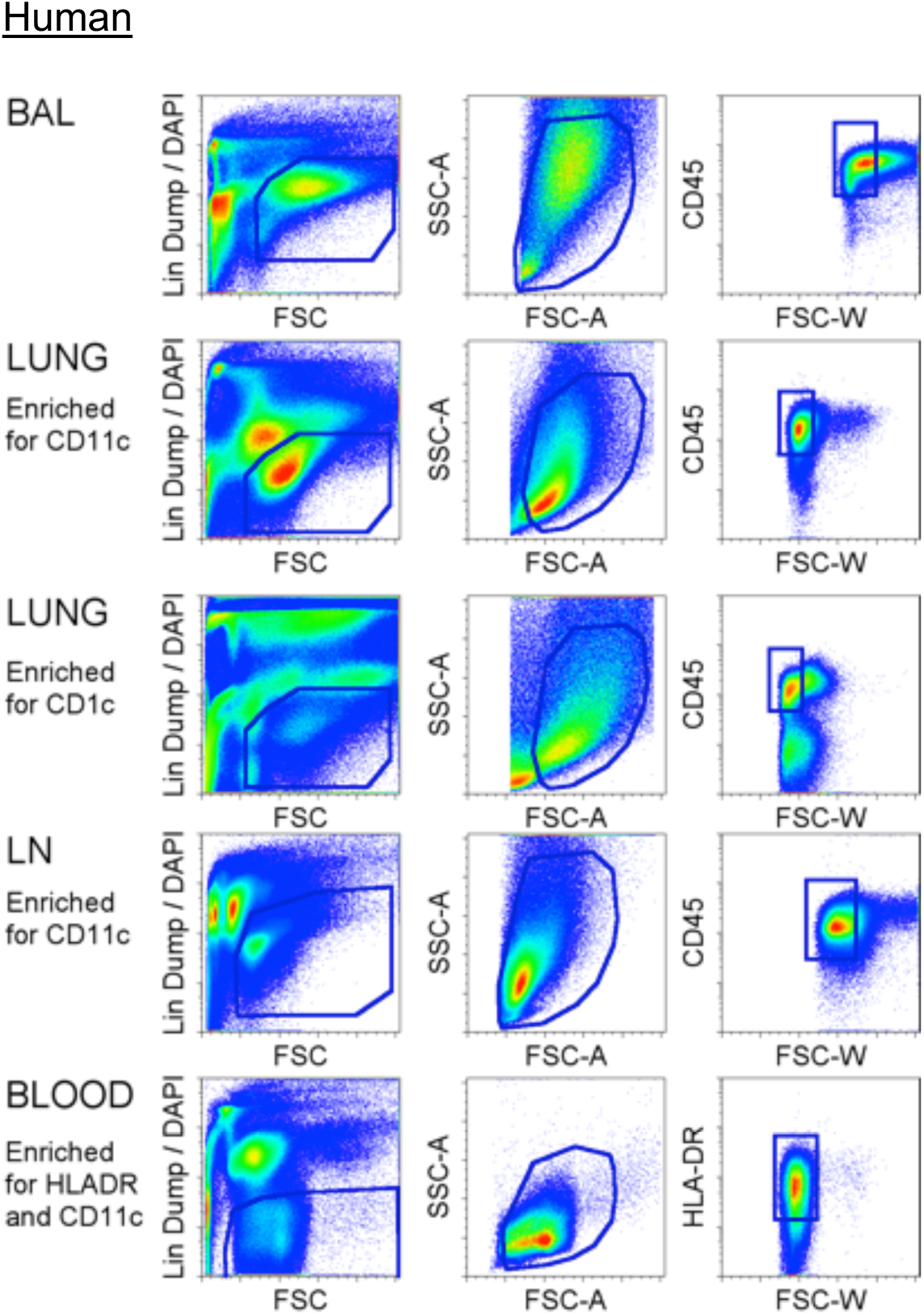
Human MP initial sorting strategy. Prior to identification of MPs, FACS gating was used to exclude Lin^+^ cells, dead cells and doublets. Lineage (Lin) dump for each tissue included antibodies against CD3, CD20, CD15 and CD56. Dying cells were labeled with 4’,6-diamidino-2-phenylindole (DAPI). Forward scatter (FSC) and side scatter (SSC) properties were used to distinguish whole cells from subcellular debris, and single cells from groups of two cells or more. Expression of CD45 was used to separate MPs from non-hematopoietic tissue cells.

**Figure S2.**
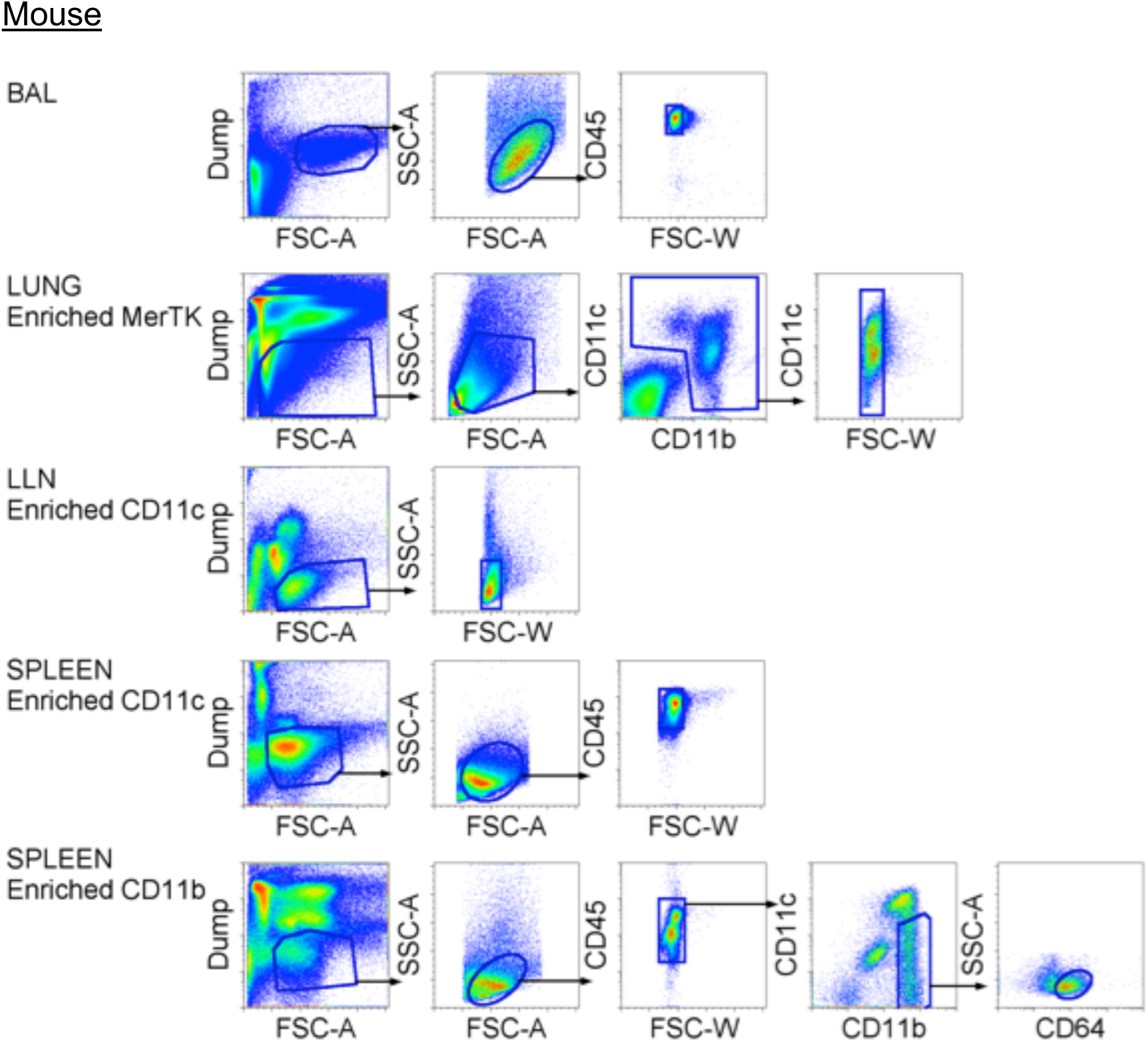
Mouse MP initial sorting strategy. Prior to identification of MPs, FACS gating was used to exclude Lin+ cells, dead cells, and doublets. Forward scatter (FSC) and side scatter (SSC) properties were used to distinguish whole cells from subcellular debris and single cells from doublets. Lineage (Lin) dump for each tissue included antibodies against CD3, B220, Ly6G and NK1.1. 4’,6-diamidino-2-phenylindole (DAPI) was used to label dying cells. For DCs, tissue-specific dump antibodies included Siglec F, CD43 and Gr1 (Lung), Gr1 and CD64 (CD11c^+^ LLN and Spleen). Additional gates confirmed hematopoietic (CD45^+^) or myeloid lineage (CD11c^+^ or CD11b^+^).

**Figure S3.**
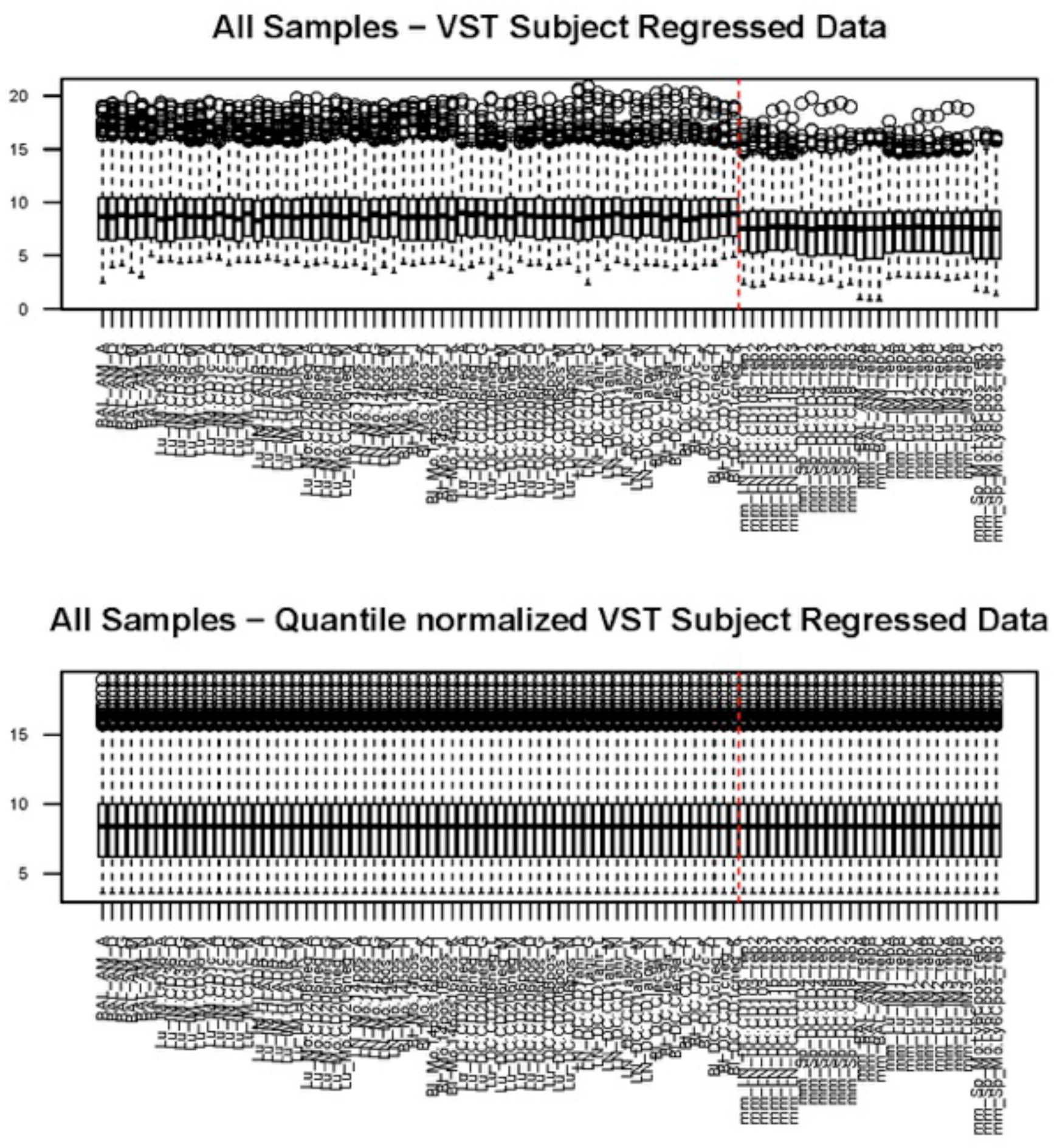
Quantile normalization of VST subject-regressed data. Expression data is shown as boxplots for the VST subject-regressed data for all samples before and after quantile normalization. A vertical red line separates human subtypes on the left from mouse subtypes on the right. Note that before quantile normalization, the mouse samples generally had lower medians than the human samples.

**Figure S4.**
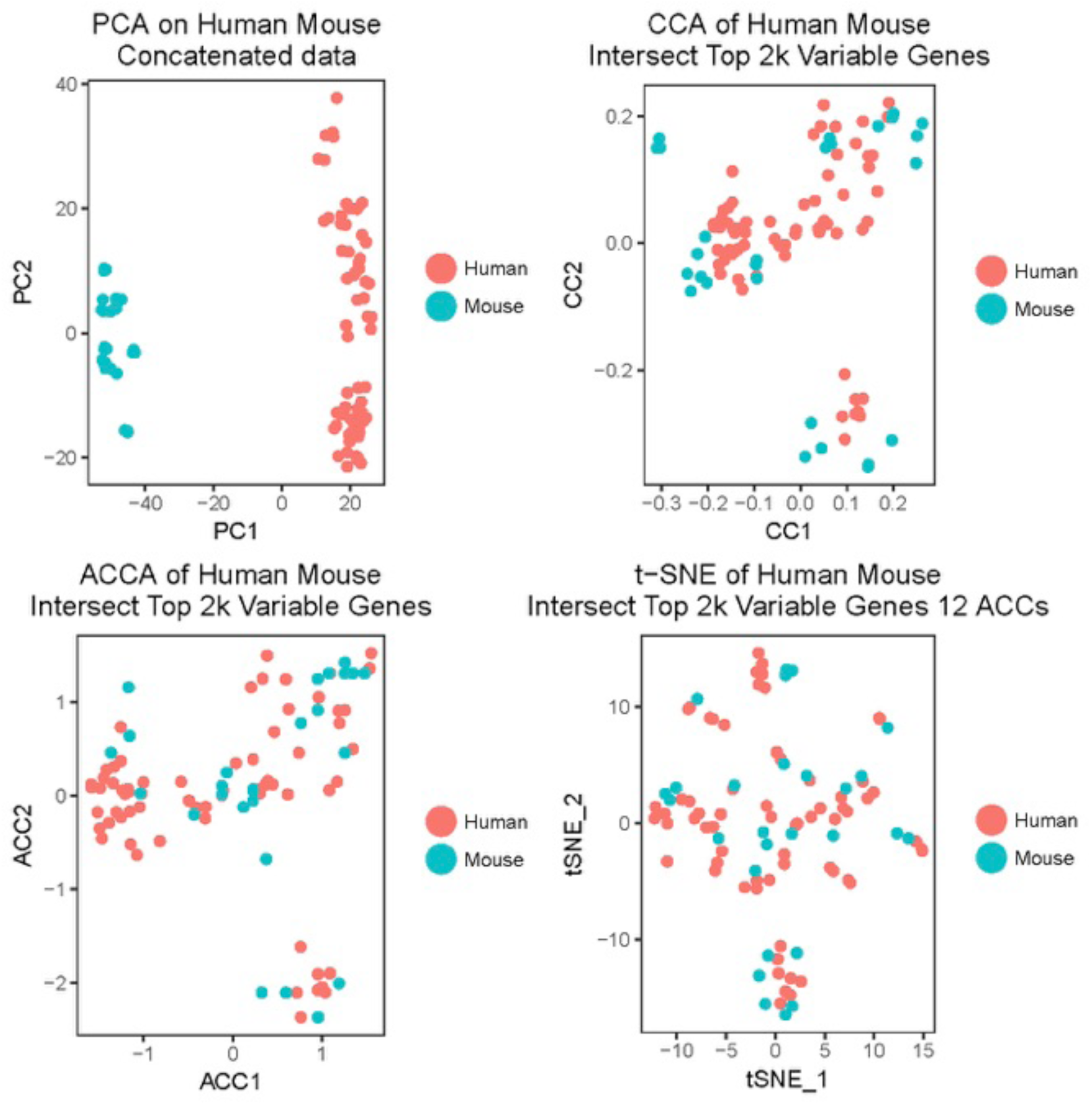
Alignment of human and mouse cell types using Seurat v2.0. Human and mouse quantile normalized VST subject-regressed expression data is subjected to the alignment method in Seurat v2. A principal component analysis (PCA) of the combined human and mouse quantile normalized data shows clear separation per species even after quantile normalization. A canonical correlation analysis (CCA) of the genes in common among the top 2000 variable genes in each species is performed and visualized as the top 2 CCs for each species. The CCs are aligned via a dynamic time-warping algorithm to bring the two CC spaces into the same coordinate space. The top two resulting aligned CCs (ACCs) are used to visualize the combined data. T-distributed Stochastic Neighbor Embedding (t-SNE) of the top 12 ACCs shows that the human and mouse data no longer cluster solely by species.

**Figure S5.**
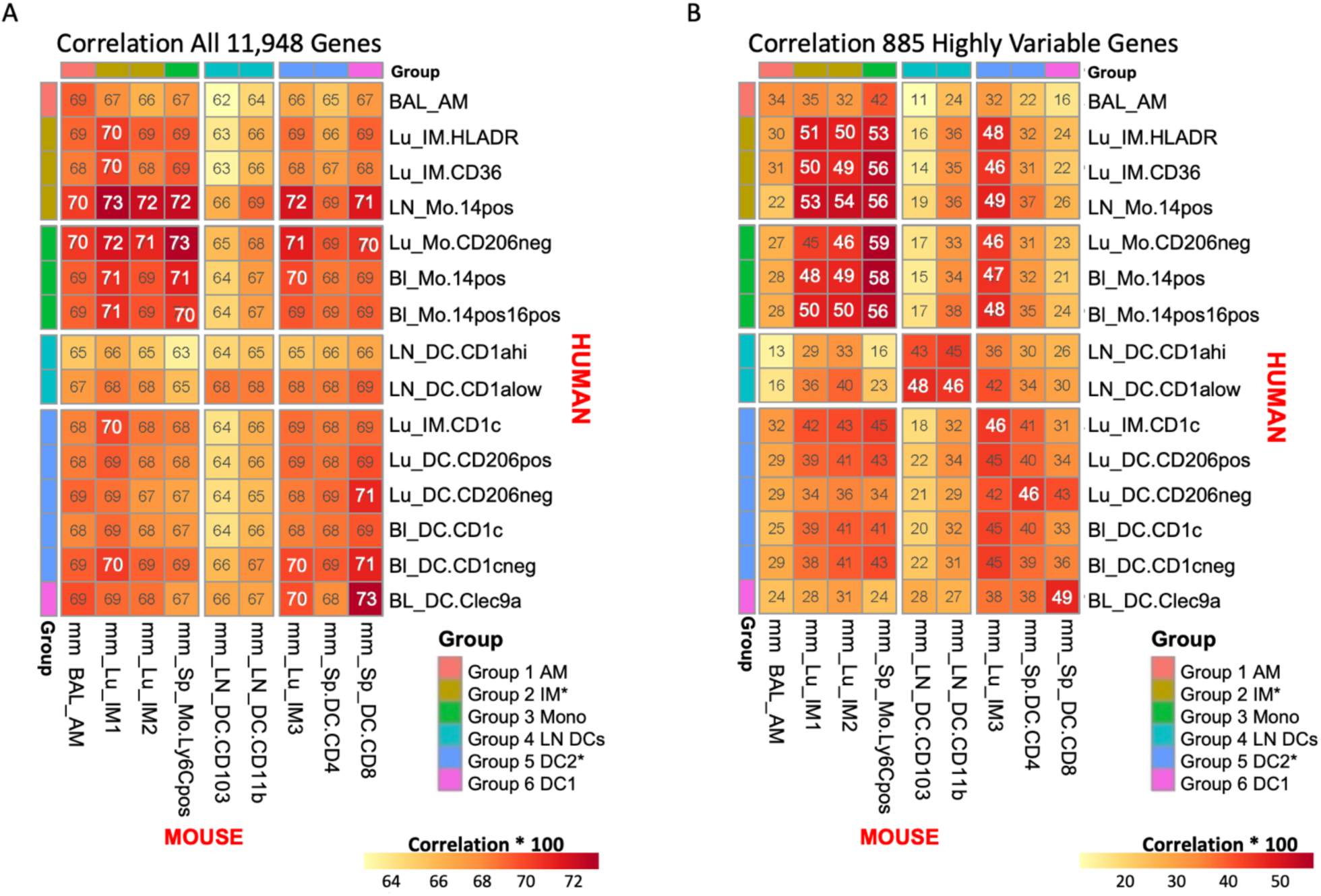
Cross-species correlation of MP subtypes. The median Pearson correlation is computed over all pairwise Pearson correlations between human samples for one cell type and mouse samples for one cell type. A) Correlation using all 11,948 autosomal homologous genes. Note that the correlation range is quite high and has a narrow range (r=64-72), such that any cell type appears equally closely related to most cross-species cell types. This may be attributed to a large fraction of all genes having low or constant expression. B) Corrrelation of the 885 most highly variable genes (hvg) shared among the top 2000 human hvgs and the top 2000 mouse hvgs. The range of correlations overall has dropped to r=20-50 but for a given cell type, multiple cross-species cell type seem nearly equally good homologs. Moreover, the best match suggested by A) does not always agree with the best match suggested by B), so determining best match by correlation is not robust to the input gene set.

**Figure S6.**
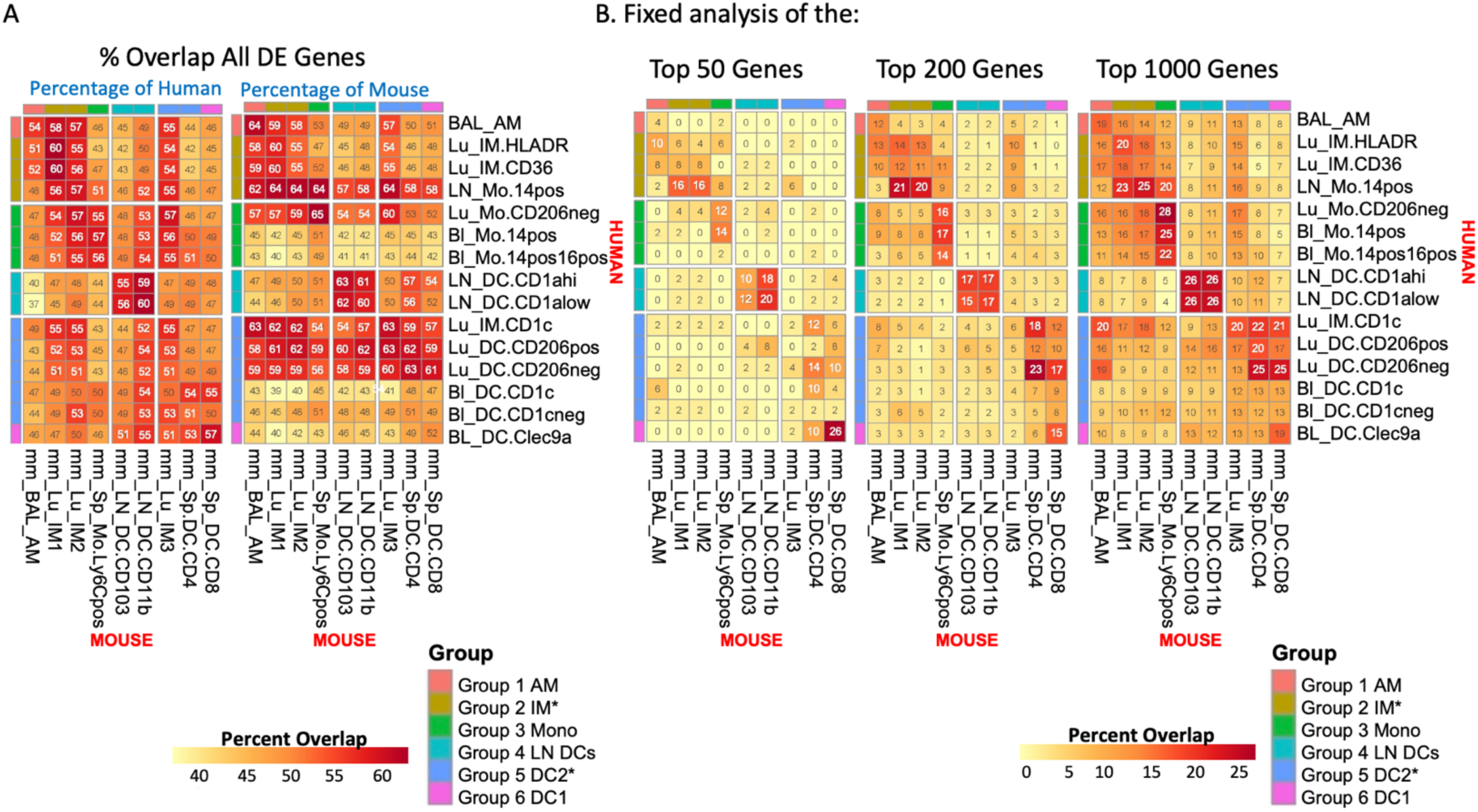
Cross-species percentage overlap of marker gene sets. A marker gene set includes genes with positive Z-score (see Methods) indicating over-expression in the cell type and p.adj > 0.05. Marker genes are ranked in decreasing order by score = [median scaled quantile normalized data]*Z-score. A) Percent overlap of all marker genes per cell type. Since the marker gene sets are not symmetric, values are shown using both human gene set sizes as the denominator for the overlap percentages and mouse. The number of marker genes ranges from approximately 4,000 to 7,000 of the original 11,948, so given the relative size of the marker gene set to the total gene size, high overlap is expected (40-68%). Note that using all DE genes is not discriminatively indicating which pairing is the best match. B) Percent overlap using fixed top marker gene sets. The percentage overlaps are symmetric given a fixed gene set size. The percent overlaps are smaller (0-25%) but are more discriminatory than using all DE genes. The maximum pairing changes given the size of the marker gene list, though several pairings are robustly suggested across all marker set sizes. This motivates the use of the average overlap percentage over the range 1-1000 as the criterion for selecting the best match.

**Figure S7.**
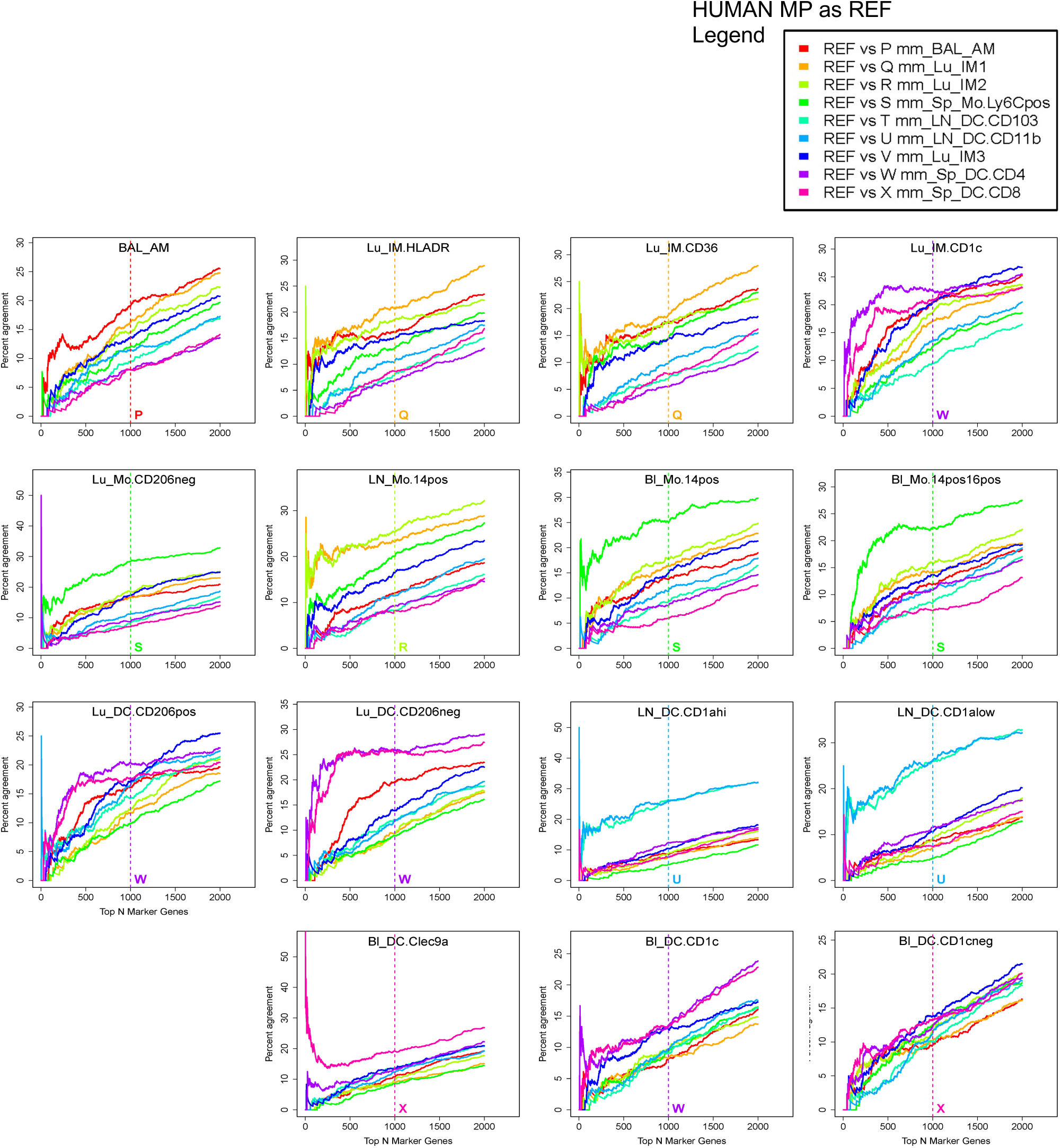
Correspondence-at-the-Top (CAT) plots in human. Candidate marker genes are ranked by their score, which multiplies their expression level scaled relative to the median expression by their marker gene Z-score (see Methods). A) For each reference MP subtype, the proportion of agreement of ranked marker gene lists versus ranked marker gene lists for each candidate subtype in the other species is calculated for progressively larger list sizes and visualized with a Correspondence-at-the-Top (CAT) plot. Lines are labeled by letters from Figure 3 indicating the subtype compared to the reference shown in the title of the figure. A vertical line indicates which MP subtype have the highest mean CAT overlap across all list sizes from 1 to 1000 with respect to the reference.

**Figure S8.**
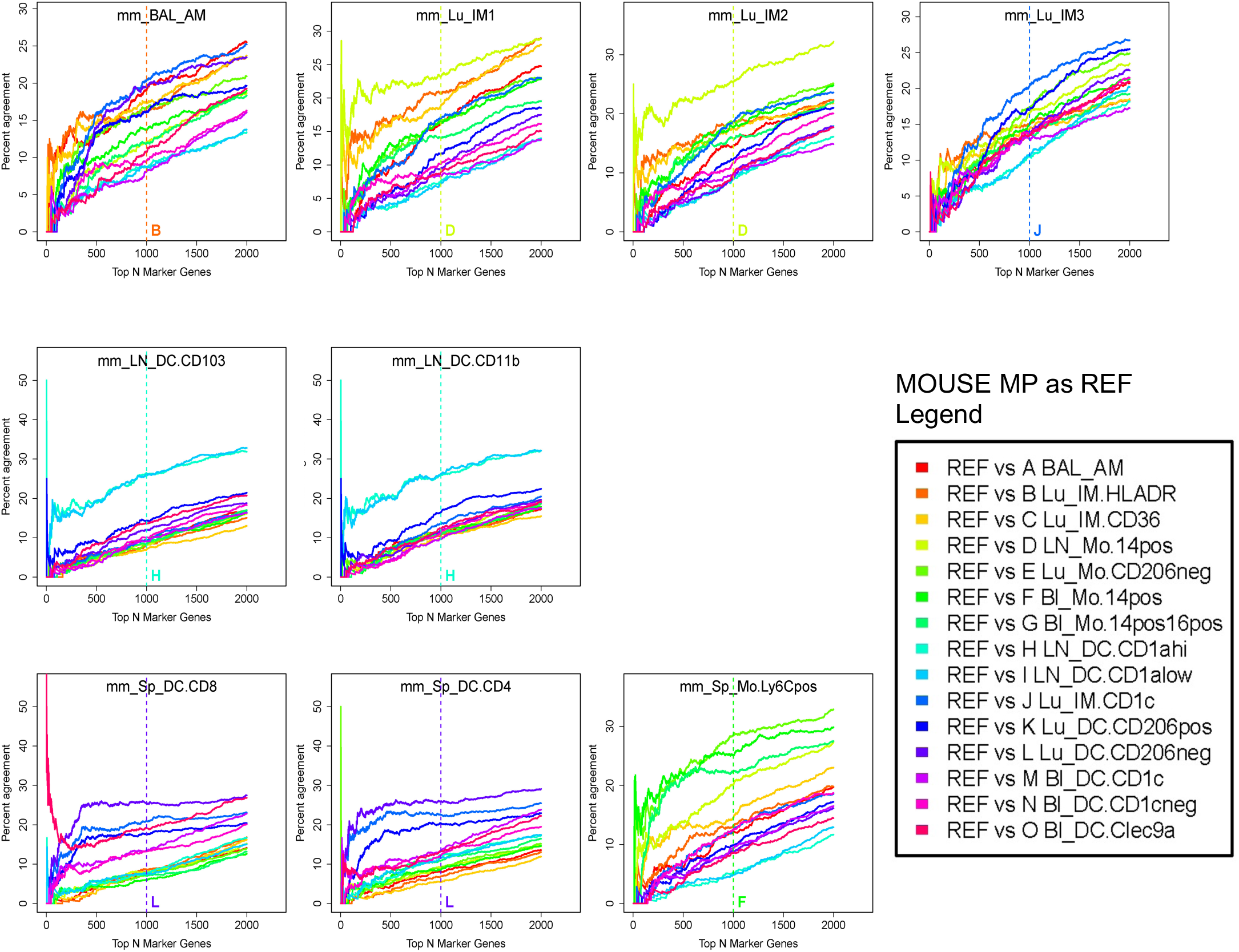
Correspondence-at-the-Top (CAT) plots in mouse. Candidate marker genes are ranked by their score, which multiplies their expression levels scaled relative to the median expression by their marker gene Z-score (see Methods). A) For each reference MP subtype, the proportion of agreement of ranked marker gene lists versus ranked marker gene lists for each candidate subtype in the other species is calculated for progressively larger list sizes and visualized with a Correspondence-at-the-Top (CAT) plot. Lines are labeled by letters from Figure 3 indicating the subtype compared to the reference shown in the title of the figure. A vertical line indicates which MP subtype had the highest mean CAT overlap across all list sizes from 1 to 1000 with respect to the reference.

**Figure S9.**
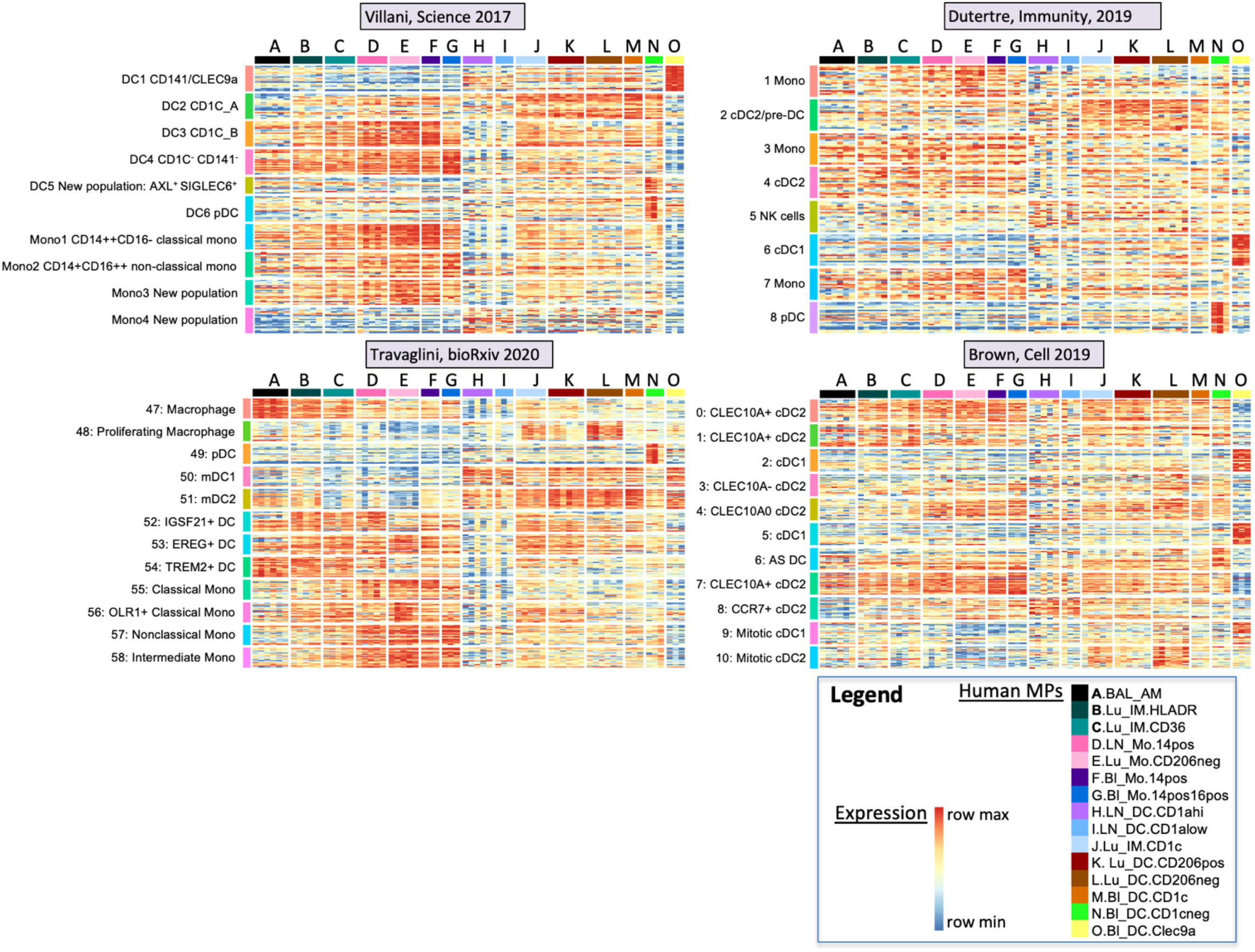
Expression of previous DC and monocyte signature genes. Gene signatures from Villani et al., (Villani et al., 2017) Dutertre et al., (Dutertre et al., 2019) Brown et al., (Brown et al., 2019) and Travaglini et al., (Travaglini https://www.biorxiv.org/content/10.1101/742320v2] are shown as heatmaps of the top 20 signature genes expressed in the 15 human subtypes, denoted by the letters A through O. The data are scaled across the row by the gene min and max to emphasize in which subtype the gene is most expressed. Correspondence of signature gene sets identified in these studies with the expected blood cell types in our MP data lends further validity to our investigation of the less well studied MP subtypes in lung and LN, as well as our cross-species MP analysis. As expected, our human blood Clec9a+ DCs corresponded most highly with the DC1 CD141/CLEC9a subset from Villani et al., (Villani et al., 2017) and the cDC1 subsets from Dutertre et al., (Dutertre et al., 2019) and Brown et al., (Brown et al., 2019). The CD1c signatures from Villani et al., were expressed in our blood DC CD1c samples. Our blood DC CD1c-cells expressed genes from the pDC signatures in Villani and Dutertre et al., papers. The blood monocytes showed high expression of Villani’s monocyte signatures: Blood monocytes CD14+ showed enrichment especially in Villani et al.’s Mono1 signature, while blood CD14+CD16+ monocytes showed enrichment in Villani et al.’s Mono2 signature. High expression of DC signature genes from Villani et al. and Brown et al., which show high expression in our monocytes, can be explained by the fact that those signatures were not derived by comparison to monocyte data. Dutertre et al. note that Villani et al.’s DC4 signature could correspond to CD16+ monocytes, which is also observed in our data. However, our LN DCs do not quite match any of the single-cell blood data sets, except perhaps Villani et al.’s Mono4 or Brown et al.’s CCR7+ cDC2 populations. The latter is interesting and ties back to the knowledge that human splenic DCs reflect a more mature signature than blood DCs, along with expressing CCR7.

**Figure S10-33. Heatmaps of top 50 genes per cell type**

The expression values are shown for the top 50 marker genes in the reference MP subtype. The subtypes are ordered by degree of overlap with the reference cell type, if within the same species, or by the mean CAT overlap shown in Figure 4B if in the other species. Samples are annotated by their cell type and tissue type. The data are scaled across the row by the gene min and max within each species to emphasize in which subtypes the gene is most expressed. The rank of the gene in the reference subtype is shown next to the gene name. Genes that also appear in the top 1000 ranked genes in the three highest corresponding subtypes in the opposing species are annotated with *, ** or ***, respectively. For example, in Figure S10, SORT1 of rank 46 in the mouse AM reference subtype is among the top 1000 marker genes in the first and third closest corresponding human subtypes, Lu IM HLADR and Lu IM CD1c, respectively. MARCO of rank 10 in the reference is among the top 1000 in all three closest subtypes. Three replicates per subtype are shown.

## References

Alferink, J., Lieberam, I., Reindl, W., Behrens, A., Weiss, S., Huser, N., Gerauer, K., Ross, R., Reske-Kunz, A. B., Ahmad-Nejad, P., Wagner, H. & Forster, I. 2003. Compartmentalized production of Ccl17 in vivo: strong inducibility in peripheral dendritic cells contrasts selective absence from the spleen. J Exp Med, 197, 585–99.

Altmann, D. M. 2018. A Nobel Prize-worthy pursuit: cancer immunology and harnessing immunity to tumour neoantigens. Immunology, 155, 283–284.

Atif, S. M., Gibbings, S. L. & Jakubzick, C. V. 2018a. Isolation and Identification of Interstitial Macrophages from the Lungs Using Different Digestion Enzymes and Staining Strategies. Methods Mol Biol, 1784, 69–76.

Atif, S. M., Gibbings, S. L., Redente, E. F., Camp, F. A., Torres, R. M., Kedl, R. M., Henson, P. M. & Jakubzick, C. V. 2018b. Immune Surveillance by Natural IgM is Required For Early Neoantigen Recognition and Initiation of Adaptive Immunity. Am J Respir Cell Mol Biol.

Atif, S. M., Nelsen, M. K., Gibbings, S. L., Desch, A. N., Kedl, R. M., Gill, R. G., Marrack, P., Murphy, K. M., Grazia, T. J., Henson, P. M. & Jakubzick, C. V. 2015. Cutting Edge: Roles for Batf3-Dependent Apcs in the Rejection of Minor Histocompatibility Antigen-Mismatched Grafts. J Immunol, 195, 46–50.

Bachem, A., Guttler, S., Hartung, E., Ebstein, F., Schaefer, M., Tannert, A., Salama, A., Movassaghi, K., Opitz, C., Mages, H. W., Henn, V., Kloetzel, P. M., Gurka, S. & Kroczek, R. A. 2010. Superior antigen cross-presentation and Xcr1 expression define human Cd11c+Cd141+ cells as homologues of mouse Cd8+ dendritic cells. J Exp Med, 207, 1273–81.

Bain, C. C., Hawley, C. A., Garner, H., Scott, C. L., Schridde, A., Steers, N. J., Mack, M., Joshi, A., Guilliams, M., Mowat, A. M., Geissmann, F. & Jenkins, S. J. 2016. Long-lived self-renewing bone marrow-derived macrophages displace embryo-derived cells to inhabit adult serous cavities. Nat Commun, 7, ncomms11852.

Bajpai, G., Schneider, C., Wong, N., Bredemeyer, A., Hulsmans, M., Nahrendorf, M., Epelman, S., Kreisel, D., Liu, Y., Itoh, A., Shankar, T. S., Selzman, C. H., Drakos, S. G. & Lavine, K. J. 2018. The human heart contains distinct macrophage subsets with divergent origins and functions. Nat Med, 24, 1234–1245.

Begley, C. G. & Ellis, L. M. 2012. Drug development: Raise standards for preclinical cancer research. Nature, 483, 531–3.

Bharat, A., Bhorade, S. M., Morales-Nebreda, L., Mc Quattie-Pimentel, A. C., Soberanes, S., Ridge, K., Decamp, M. M., Mestan, K. K., Perlman, H., Budinger, G. R. & Misharin, A. V. 2015. Flow Cytometry Reveals Similarities Between Lung Macrophages in Humans and Mice. Am J Respir Cell Mol Biol.

Bharat, A., Bhorade, S. M., Morales-Nebreda, L., Mcquattie-Pimentel, A. C., Soberanes, S., Ridge, K., Decamp, M. M., Mestan, K. K., Perlman, H., Budinger, G. R. & Misharin, A. V. 2016. Flow Cytometry Reveals Similarities Between Lung Macrophages in Humans and Mice. Am J Respir Cell Mol Biol, 54, 147–9.

Bolstad, B. M., Irizarry, R. A., Astrand, M. & Speed, T. P. 2003. A comparison of normalization methods for high density oligonucleotide array data based on variance and bias. Bioinformatics, 19, 185–93.

Bosteels, C., Neyt, K., Vanheerswynghels, M., Van Helden, M. J., Sichien, D., Debeuf, N., De Prijck, S., Bosteels, V., Vandamme, N., Martens, L., Saeys, Y., Louagie, E., Lesage, M., Williams, D. L., Tang, S. C., Mayer, J. U., Ronchese, F., Scott, C. L., Hammad, H., Guilliams, M. & Lambrecht, B. N. 2020. Inflammatory Type 2 cdcs Acquire Features of cdc1s and Macrophages to Orchestrate Immunity to Respiratory Virus Infection. Immunity, 52, 1039–1056 e9.

Brown, C. C., Gudjonson, H., Pritykin, Y., Deep, D., Lavallee, V. P., Mendoza, A., Fromme, R., Mazutis, L., Ariyan, C., Leslie, C., Pe’er, D. & Rudensky, A. Y. 2019. Transcriptional Basis of Mouse and Human Dendritic Cell Heterogeneity. Cell, 179, 846–863 e24.

Butler, A., Hoffman, P., Smibert, P., Papalexi, E. & Satija, R. 2018. Integrating single-cell transcriptomic data across different conditions, technologies, and species. Nat Biotechnol, 36, 411–420.

Caminschi, I., Proietto, A. I., Ahmet, F., Kitsoulis, S., Shin Teh, J., Lo, J. C., Rizzitelli, A., Wu, L., Vremec, D., Van Dommelen, S. L., Campbell, I. K., Maraskovsky, E., Braley, H., Davey, G. M., Mottram, P., Van De Velde, N., Jensen, K., Lew, A. M., Wright, M. D., Heath, W. R., Shortman, K. & Lahoud, M. H. 2008. The dendritic cell subtype-restricted C-type lectin Clec9A is a target for vaccine enhancement. Blood, 112, 3264–73.

Chakarov, S., Lim, H. Y., Tan, L., Lim, S. Y., See, P., Lum, J., Zhang, X. M., Foo, S., Nakamizo, S., Duan, K., Kong, W. T., Gentek, R., Balachander, A., Carbajo, D., Bleriot, C., Malleret, B., Tam, J. K. C., Baig, S., Shabeer, M., Toh, S. E. S., Schlitzer, A., Larbi, A., Marichal, T., Malissen, B., Chen, J., Poidinger, M., Kabashima, K., Bajenoff, M., Ng, L. G., Angeli, V. & Ginhoux, F. 2019. Two distinct interstitial macrophage populations coexist across tissues in specific subtissular niches. Science, 363.

Desch, A. N., Gibbings, S. L., Clambey, E. T., Janssen, W. J., Slansky, J. E., Kedl, R. M., Henson, P. M. & Jakubzick, C. 2014. Dendritic cell subsets require cis-activation for cytotoxic Cd8 T-cell induction. Nat Commun, 5, 4674.

Desch, A. N., Gibbings, S. L., Goyal, R., Kolde, R., Bednarek, J., Bruno, T., Slansky, J. E., Jacobelli, J., Mason, R., Ito, Y., Messier, E., Randolph, G. J., Prabagar, M., Atif, S. M., Segura, E., Xavier, R. J., Bratton, D. L., Janssen, W. J., Henson, P. M. & Jakubzick, C. V. 2015. Flow Cytometric Analysis of Mononuclear Phagocytes in Non-diseased Human Lung and Lung-draining Lymph Nodes. Am J Respir Crit Care Med.

Desch, A. N., Randolph, G. J., Murphy, K., Gautier, E. L., Kedl, R. M., Lahoud, M. H., Caminschi, I., Shortman, K., Henson, P. M. & Jakubzick, C. V. 2011. Cd103+ pulmonary dendritic cells preferentially acquire and present apoptotic cell-associated antigen. J Exp Med, 208, 1789–97.

Dobin, A., Davis, C. A., Schlesinger, F., Drenkow, J., Zaleski, C., Jha, S., Batut, P., Chaisson, M. & Gingeras, T. R. 2013. Star: ultrafast universal Rna-seq aligner. Bioinformatics, 29, 15–21.

Dutertre, C. A., Becht, E., Irac, S. E., Khalilnezhad, A., Narang, V., Khalilnezhad, S., Ng, P. Y., Van Den Hoogen, L. L., Leong, J. Y., Lee, B., Chevrier, M., Zhang, X. M., Yong, P. J. A., Koh, G., Lum, J., Howland, S. W., Mok, E., Chen, J., Larbi, A., Tan, H. K. K., Lim, T. K. H., Karagianni, P., Tzioufas, A. G., Malleret, B., Brody, J., Albani, S., Van Roon, J., Radstake, T., Newell, E. W. & Ginhoux, F. 2019. Single-Cell Analysis of Human Mononuclear Phagocytes Reveals Subset-Defining Markers and Identifies Circulating Inflammatory Dendritic Cells. Immunity, 51, 573–589 e8.

Epelman, S., Lavine, K. J., Beaudin, A. E., Sojka, D. K., Carrero, J. A., Calderon, B., Brija, T., Gautier, E. L., Ivanov, S., Satpathy, A. T., Schilling, J. D., Schwendener, R., Sergin, I., Razani, B., Forsberg, E. C., Yokoyama, W. M., Unanue, E. R., Colonna, M., Randolph, G. J. & Mann, D. L. 2014. Embryonic and adult-derived resident cardiac macrophages are maintained through distinct mechanisms at steady state and during inflammation. Immunity, 40, 91–104.

Forster, R., Schubel, A., Breitfeld, D., Kremmer, E., Renner-Muller, I., Wolf, E. & Lipp, M. 1999. Ccr7 coordinates the primary immune response by establishing functional microenvironments in secondary lymphoid organs. Cell, 99, 23–33.

Gautier, E. L., Shay, T., Miller, J., Greter, M., Jakubzick, C., Ivanov, S., Helft, J., Chow, A., Elpek, K. G., Gordonov, S., Mazloom, A. R., Ma’ayan, A., Chua, W. J., Hansen, T. H., Turley, S. J., Merad, M., Randolph, G. J. & Immunological Genome, C. 2012. Gene-expression profiles and transcriptional regulatory pathways that underlie the identity and diversity of mouse tissue macrophages. Nat Immunol, 13, 1118–28.

Gibbings, S. L. & Jakubzick, C. V. 2018a. A Consistent Method to Identify and Isolate Mononuclear Phagocytes from Human Lung and Lymph Nodes. Methods Mol Biol, 1799, 381–395.

Gibbings, S. L. & Jakubzick, C. V. 2018b. Isolation and Characterization of Mononuclear Phagocytes in the Mouse Lung and Lymph Nodes. Methods Mol Biol, 1809, 33–44.

Gibbings, S. L., Thomas, S. M., Atif, S. M., Mccubbrey, A. L., Desch, A. N., Danhorn, T., Leach, S. M., Bratton, D. L., Henson, P. M., Janssen, W. J. & Jakubzick, C. V. 2017. Three Unique Interstitial Macrophages in the Murine Lung at Steady State. Am J Respir Cell Mol Biol, 57, 66–76.

Guilliams, M., Dutertre, C. A., Scott, C. L., Mcgovern, N., Sichien, D., Chakarov, S., Van Gassen, S., Chen, J., Poidinger, M., De Prijck, S., Tavernier, S. J., Low, I., Irac, S. E., Mattar, C. N., Sumatoh, H. R., Low, G. H. L., Chung, T. J. K., Chan, D. K. H., Tan, K. K., Hon, T. L. K., Fossum, E., Bogen, B., Choolani, M., Chan, J. K. Y., Larbi, A., Luche, H., Henri, S., Saeys, Y., Newell, E. W., Lambrecht, B. N., Malissen, B. & Ginhoux, F. 2016. Unsupervised High-Dimensional Analysis Aligns Dendritic Cells across Tissues and Species. Immunity, 45, 669–684.

Guilliams, M., Lambrecht, B. N. & Hammad, H. 2013. Division of labor between lung dendritic cells and macrophages in the defense against pulmonary infections. Mucosal Immunol, 6, 464–73.

Guilliams, M. & Scott, C. L. 2017. Does niche competition determine the origin of tissue-resident macrophages? Nat Rev Immunol, 17, 451–460.

Haniffa, M., Shin, A., Bigley, V., Mcgovern, N., Teo, P., See, P., Wasan, P. S., Wang, X. N., Malinarich, F., Malleret, B., Larbi, A., Tan, P., Zhao, H., Poidinger, M., Pagan, S., Cookson, S., Dickinson, R., Dimmick, I., Jarrett, R. F., Renia, L., Tam, J., Song, C., Connolly, J., Chan, J. K., Gehring, A., Bertoletti, A., Collin, M. & Ginhoux, F. 2012. Human tissues contain Cd141hi cross-presenting dendritic cells with functional homology to mouse Cd103+ nonlymphoid dendritic cells. Immunity, 37, 60–73.

Hildner, K., Edelson, B. T., Purtha, W. E., Diamond, M., Matsushita, H., Kohyama, M., Calderon, B., Schraml, B. U., Unanue, E. R., Diamond, M. S., Schreiber, R. D., Murphy, T. L. & Murphy, K. M. 2008. Batf3 deficiency reveals a critical role for Cd8alpha+ dendritic cells in cytotoxic T cell immunity. Science, 322, 1097–100.

Huysamen, C., Willment, J. A., Dennehy, K. M. & Brown, G. D. 2008. Clec9A is a novel activation C-type lectin-like receptor expressed on Bdca3+ dendritic cells and a subset of monocytes. J Biol Chem, 283, 16693–701.

Ingersoll, M. A., Spanbroek, R., Lottaz, C., Gautier, E. L., Frankenberger, M., Hoffmann, R., Lang, R., Haniffa, M., Collin, M., Tacke, F., Habenicht, A. J., Ziegler-Heitbrock, L. & Randolph, G. J. 2010. Comparison of gene expression profiles between human and mouse monocyte subsets. Blood, 115, e10–9.

Irizarry, R. A., Warren, D., Spencer, F., Kim, I. F., Biswal, S., Frank, B. C., Gabrielson, E., Garcia, J. G., Geoghegan, J., Germino, G., Griffin, C., Hilmer, S. C., Hoffman, E., Jedlicka, A. E., Kawasaki, E., Martinez-Murillo, F., Morsberger, L., Lee, H., Petersen, D., Quackenbush, J., Scott, A., Wilson, M., Yang, Y., Ye, S. Q. & Yu, W. 2005. Multiple-laboratory comparison of microarray platforms. Nat Methods, 2, 345–50.

Jakubzick, C., Gautier, E. L., Gibbings, S. L., Sojka, D. K., Schlitzer, A., Johnson, T. E., Ivanov, S., Duan, Q., Bala, S., Condon, T., Van Rooijen, N., Grainger, J. R., Belkaid, Y., Ma’ayan, A., Riches, D. W., Yokoyama, W. M., Ginhoux, F., Henson, P. M. & Randolph, G. J. 2013. Minimal differentiation of classical monocytes as they survey steady-state tissues and transport antigen to lymph nodes. Immunity, 39, 599–610.

Jakubzick, C., Helft, J., Kaplan, T. J. & Randolph, G. J. 2008. Optimization of methods to study pulmonary dendritic cell migration reveals distinct capacities of Dc subsets to acquire soluble versus particulate antigen. J Immunol Methods, 337, 121–31.

Jakubzick, C., Tacke, F., Llodra, J., Van Rooijen, N. & Randolph, G. J. 2006. Modulation of dendritic cell trafficking to and from the airways. J Immunol, 176, 3578–84.

Jakubzick, C. V., Randolph, G. J. & Henson, P. M. 2017. Monocyte differentiation and antigen-presenting functions. Nat Rev Immunol, 17, 349–362.

Jiang, H., Lei, R., Ding, S. W. & Zhu, S. 2014. Skewer: a fast and accurate adapter trimmer for next-generation sequencing paired-end reads. Bmc Bioinformatics, 15, 182.

Jongbloed, S. L., Kassianos, A. J., Mcdonald, K. J., Clark, G. J., Ju, X., Angel, C. E., Chen, C. J., Dunbar, P. R., Wadley, R. B., Jeet, V., Vulink, A. J., Hart, D. N. & Radford, K. J. 2010. Human Cd141+ (Bdca-3)+ dendritic cells (Dcs) represent a unique myeloid Dc subset that cross-presents necrotic cell antigens. J Exp Med, 207, 1247–60.

Kim, T. S., Gorski, S. A., Hahn, S., Murphy, K. M. & Braciale, T. J. 2014. Distinct dendritic cell subsets dictate the fate decision between effector and memory Cd8(+) T cell differentiation by a Cd24-dependent mechanism. Immunity, 40, 400–13.

Kuznetsova, A., Brockhoff, P. B. & Christensen, R. H. B. 2017. lmerTest Package: Tests in Linear Mixed Effects Models. Journal of Statistical Software, 25418.

Larson, S. R., Atif, S. M., Gibbings, S. L., Thomas, S. M., Prabagar, M. G., Danhorn, T., Leach, S. M., Henson, P. M. & Jakubzick, C. V. 2016. Ly6C(+) monocyte efferocytosis and cross-presentation of cell-associated antigens. Cell Death Differ, 23, 997–1003.

Leon, B. & Ardavin, C. 2008. Monocyte migration to inflamed skin and lymph nodes is differentially controlled by L-selectin and Psgl-1. Blood, 111, 3126–30.

Liao, Y., Smyth, G. K. & Shi, W. 2019. The R package Rsubread is easier, faster, cheaper and better for alignment and quantification of Rna sequencing reads. Nucleic Acids Res, 47, e47.

Lim, H. Y., Lim, S. Y., Tan, C. K., Thiam, C. H., Goh, C. C., Carbajo, D., Chew, S. H. S., See, P., Chakarov, S., Wang, X. N., Lim, L. H., Johnson, L. A., Lum, J., Fong, C. Y., Bongso, A., Biswas, A., Goh, C., Evrard, M., Yeo, K. P., Basu, R., Wang, J. K., Tan, Y., Jain, R., Tikoo, S., Choong, C., Weninger, W., Poidinger, M., Stanley, R. E., Collin, M., Tan, N. S., Ng, L. G., Jackson, D. G., Ginhoux, F. & Angeli, V. 2018. Hyaluronan Receptor Lyve-1-Expressing Macrophages Maintain Arterial Tone through Hyaluronan-Mediated Regulation of Smooth Muscle Cell Collagen. Immunity, 49, 326–341 e7.

Liu, K., Victora, G. D., Schwickert, T. A., Guermonprez, P., Meredith, M. M., Yao, K., Chu, F. F., Randolph, G. J., Rudensky, A. Y. & Nussenzweig, M. 2009. In vivo analysis of dendritic cell development and homeostasis. Science, 324, 392–7.

Love, M. I., Huber, W. & Anders, S. 2014. Moderated estimation of fold change and dispersion for Rna-seq data with Deseq2. Genome Biol, 15, 550.

Masopust, D., Sivula, C. P. & Jameson, S. C. 2017. Of Mice, Dirty Mice, and Men: Using Mice To Understand Human Immunology. J Immunol, 199, 383–388.

Mccubbrey, A. L., Barthel, L., Mohning, M. P., Redente, E. F., Mould, K. J., Thomas, S. M., Leach, S. M., Danhorn, T., Gibbings, S. L., Jakubzick, C. V., Henson, P. M. & Janssen, W. J. 2018. Deletion of c-Flip from Cd11b(hi) Macrophages Prevents Development of Bleomycin-induced Lung Fibrosis. Am J Respir Cell Mol Biol, 58, 66–78.

Misharin, A. V., Morales-Nebreda, L., Mutlu, G. M., Budinger, G. R. & Perlman, H. 2013. Flow cytometric analysis of macrophages and dendritic cell subsets in the mouse lung. Am J Respir Cell Mol Biol, 49, 503–10.

Misharin, A. V., Morales-Nebreda, L., Reyfman, P. A., Cuda, C. M., Walter, J. M., Mcquattie-Pimentel, A. C., Chen, C. I., Anekalla, K. R., Joshi, N., Williams, K. J. N., Abdala-Valencia, H., Yacoub, T. J., Chi, M., Chiu, S., Gonzalez-Gonzalez, F. J., Gates, K., Lam, A. P., Nicholson, T. T., Homan, P. J., Soberanes, S., Dominguez, S., Morgan, V. K., Saber, R., Shaffer, A., Hinchcliff, M., Marshall, S. A., Bharat, A., Berdnikovs, S., Bhorade, S. M., Bartom, E. T., Morimoto, R. I., Balch, W. E., Sznajder, J. I., Chandel, N. S., Mutlu, G. M., Jain, M., Gottardi, C. J., Singer, B. D., Ridge, K. M., Bagheri, N., Shilatifard, A., Budinger, G. R. S. & Perlman, H. 2017. Monocyte-derived alveolar macrophages drive lung fibrosis and persist in the lung over the life span. J Exp Med, 214, 2387–2404.

Mould, K. J., Barthel, L., Mohning, M. P., Thomas, S. M., Mccubbrey, A. L., Danhorn, T., Leach, S. M., Fingerlin, T. E., O’connor, B. P., Reisz, J. A., D’alessandro, A., Bratton, D. L., Jakubzick, C. V. & Janssen, W. J. 2017. Cell Origin Dictates Programming of Resident versus Recruited Macrophages during Acute Lung Injury. Am J Respir Cell Mol Biol, 57, 294–306.

Mould, K. J., Jackson, N. D., Henson, P. M., Seibold, M. & Janssen, W. J. 2019. Single cell Rna sequencing identifies unique inflammatory airspace macrophage subsets. Jci Insight, 4.

Murphy, K. M. 2013. Transcriptional control of dendritic cell development. Adv Immunol, 120, 239–67.

Nakano, H., Moran, T. P., Nakano, K., Gerrish, K. E., Bortner, C. D. & Cook, D. N. 2015. Complement receptor C5aR1/Cd88 and dipeptidyl peptidase-4/Cd26 define distinct hematopoietic lineages of dendritic cells. J Immunol, 194, 3808–19.

Plantinga, M., Guilliams, M., Vanheerswynghels, M., Deswarte, K., Branco-Madeira, F., Toussaint, W., Vanhoutte, L., Neyt, K., Killeen, N., Malissen, B., Hammad, H. & Lambrecht, B. N. 2013. Conventional and monocyte-derived Cd11b(+) dendritic cells initiate and maintain T helper 2 cell-mediated immunity to house dust mite allergen. Immunity, 38, 322–35.

Randolph, G. J., Angeli, V. & Swartz, M. A. 2005. Dendritic-cell trafficking to lymph nodes through lymphatic vessels. Nat Rev Immunol, 5, 617–28.

Rapp, M., Wintergerst, M. W. M., Kunz, W. G., Vetter, V. K., Knott, M. M. L., Lisowski, D., Haubner, S., Moder, S., Thaler, R., Eiber, S., Meyer, B., Rohrle, N., Piseddu, I., Grassmann, S., Layritz, P., Kuhnemuth, B., Stutte, S., Bourquin, C., Von Andrian, U. H., Endres, S. & Anz, D. 2019. Ccl22 controls immunity by promoting regulatory T cell communication with dendritic cells in lymph nodes. J Exp Med, 216, 1170–1181.

Reyfman, P. A., Walter, J. M., Joshi, N., Anekalla, K. R., Mcquattie-Pimentel, A. C., Chiu, S., Fernandez, R., Akbarpour, M., Chen, C. I., Ren, Z., Verma, R., Abdala-Valencia, H., Nam, K., Chi, M., Han, S., Gonzalez-Gonzalez, F. J., Soberanes, S., Watanabe, S., Williams, K. J. N., Flozak, A. S., Nicholson, T. T., Morgan, V. K., Winter, D. R., Hinchcliff, M., Hrusch, C. L., Guzy, R. D., Bonham, C. A., Sperling, A. I., Bag, R., Hamanaka, R. B., Mutlu, G. M., Yeldandi, A. V., Marshall, S. A., Shilatifard, A., Amaral, L. A. N., Perlman, H., Sznajder, J. I., Argento, A. C., Gillespie, C. T., Dematte, J., Jain, M., Singer, B. D., Ridge, K. M., Lam, A. P., Bharat, A., Bhorade, S. M., Gottardi, C. J., Budinger, G. R. S. & Misharin, A. V. 2018. Single-Cell Transcriptomic Analysis of Human Lung Provides Insights into the Pathobiology of Pulmonary Fibrosis. Am J Respir Crit Care Med.

Reynolds, G. & Haniffa, M. 2015. Human and Mouse Mononuclear Phagocyte Networks: A Tale of Two Species? Front Immunol, 6, 330.

Rosenthal, R. 1978. Combining Results of Independent Studies. Psychological Bulletin, 85, 185–193.

Satija, R., Farrell, J. A., Gennert, D., Schier, A. F. & Regev, A. 2015. Spatial reconstruction of single-cell gene expression data. Nat Biotechnol, 33, 495–502.

Satpathy, A. T., Kc, W., Albring, J. C., Edelson, B. T., Kretzer, N. M., Bhattacharya, D., Murphy, T. L. & Murphy, K. M. 2012. Zbtb46 expression distinguishes classical dendritic cells and their committed progenitors from other immune lineages. J Exp Med, 209, 1135–52.

Schlitzer, A., Mcgovern, N., Teo, P., Zelante, T., Atarashi, K., Low, D., Ho, A. W., See, P., Shin, A., Wasan, P. S., Hoeffel, G., Malleret, B., Heiseke, A., Chew, S., Jardine, L., Purvis, H. A., Hilkens, C. M., Tam, J., Poidinger, M., Stanley, E. R., Krug, A. B., Renia, L., Sivasankar, B., Ng, L. G., Collin, M., Ricciardi-Castagnoli, P., Honda, K., Haniffa, M. & Ginhoux, F. 2013. Irf4 transcription factor-dependent Cd11b+ dendritic cells in human and mouse control mucosal Il-17 cytokine responses. Immunity, 38, 970–83.

Schyns, J., Bai, Q., Ruscitti, C., Radermecker, C., De Schepper, S., Chakarov, S., Farnir, F., Pirottin, D., Ginhoux, F., Boeckxstaens, G., Bureau, F. & Marichal, T. 2019. Non-classical tissue monocytes and two functionally distinct populations of interstitial macrophages populate the mouse lung. Nat Commun, 10, 3964.

Scott, C. L., Henri, S. & Guilliams, M. 2014. Mononuclear phagocytes of the intestine, the skin, and the lung. Immunol Rev, 262, 9–24.

Stutte, S., Quast, T., Gerbitzki, N., Savinko, T., Novak, N., Reifenberger, J., Homey, B., Kolanus, W., Alenius, H. & Forster, I. 2010. Requirement of Ccl17 for Ccr7- and Cxcr4-dependent migration of cutaneous dendritic cells. Proc Natl Acad Sci U S A, 107, 8736–41.

Tamoutounour, S., Guilliams, M., Montanana Sanchis, F., Liu, H., Terhorst, D., Malosse, C., Pollet, E., Ardouin, L., Luche, H., Sanchez, C., Dalod, M., Malissen, B. & Henri, S. 2013. Origins and functional specialization of macrophages and of conventional and monocyte-derived dendritic cells in mouse skin. Immunity, 39, 925–38.

Tamoutounour, S., Henri, S., Lelouard, H., De Bovis, B., De Haar, C., Van Der Woude, C. J., Woltman, A. M., Reyal, Y., Bonnet, D., Sichien, D., Bain, C. C., Mowat, A. M., Reis E Sousa, C., Poulin, L. F., Malissen, B. & Guilliams, M. 2012. Cd64 distinguishes macrophages from dendritic cells in the gut and reveals the Th1-inducing role of mesenteric lymph node macrophages during colitis. Eur J Immunol, 42, 3150–66.

Theisen, D. J., Davidson, J. T. T., Briseno, C. G., Gargaro, M., Lauron, E. J., Wang, Q., Desai, P., Durai, V., Bagadia, P., Brickner, J. R., Beatty, W. L., Virgin, H. W., Gillanders, W. E., Mosammaparast, N., Diamond, M. S., Sibley, L. D., Yokoyama, W., Schreiber, R. D., Murphy, T. L. & Murphy, K. M. 2018. Wdfy4 is required for cross-presentation in response to viral and tumor antigens. Science, 362, 694–699.

Trapnell, B. C., Carey, B. C., Uchida, K. & Suzuki, T. 2009. Pulmonary alveolar proteinosis, a primary immunodeficiency of impaired Gm-Csf stimulation of macrophages. Curr Opin Immunol, 21, 514–21.

Tussiwand, R., Everts, B., Grajales-Reyes, G. E., Kretzer, N. M., Iwata, A., Bagaitkar, J., Wu, X., Wong, R., Anderson, D. A., Murphy, T. L., Pearce, E. J. & Murphy, K. M. 2015. Klf4 expression in conventional dendritic cells is required for T helper 2 cell responses. Immunity, 42, 916–28.

Tussiwand, R., Lee, W. L., Murphy, T. L., Mashayekhi, M., Wumesh, K. C., Albring, J. C., Satpathy, A. T., Rotondo, J. A., Edelson, B. T., Kretzer, N. M., Wu, X., Weiss, L. A., Glasmacher, E., Li, P., Liao, W., Behnke, M., Lam, S. S., Aurthur, C. T., Leonard, W. J., Singh, H., Stallings, C. L., Sibley, L. D., Schreiber, R. D. & Murphy, K. M. 2012. Compensatory dendritic cell development mediated by Batf-Irf interactions. Nature, 490, 502–7.

Ural, B. B., Yeung, S. T., Damani-Yokota, P., Devlin, J. C., De Vries, M., Vera-Licona, P., Samji, T., Sawai, C. M., Jang, G., Perez, O. A., Pham, Q., Maher, L., Loke, P., Dittmann, M., Reizis, B. & Khanna, K. M. 2020. Identification of a nerve-associated, lung-resident interstitial macrophage subset with distinct localization and immunoregulatory properties. Sci Immunol, 5.

Vermaelen, K. Y., Carro-Muino, I., Lambrecht, B. N. & Pauwels, R. A. 2001. Specific migratory dendritic cells rapidly transport antigen from the airways to the thoracic lymph nodes. J Exp Med, 193, 51–60.

Villani, A. C., Satija, R., Reynolds, G., Sarkizova, S., Shekhar, K., Fletcher, J., Griesbeck, M., Butler, A., Zheng, S., Lazo, S., Jardine, L., Dixon, D., Stephenson, E., Nilsson, E., Grundberg, I., Mcdonald, D., Filby, A., Li, W., De Jager, P. L., Rozenblatt-Rosen, O., Lane, A. A., Haniffa, M., Regev, A. & Hacohen, N. 2017. Single-cell Rna-seq reveals new types of human blood dendritic cells, monocytes, and progenitors. Science, 356.

Vulcano, M., Albanesi, C., Stoppacciaro, A., Bagnati, R., D’amico, G., Struyf, S., Transidico, P., Bonecchi, R., Del Prete, A., Allavena, P., Ruco, L. P., Chiabrando, C., Girolomoni, G., Mantovani, A. & Sozzani, S. 2001. Dendritic cells as a major source of macrophage-derived chemokine/Ccl22 in vitro and in vivo. Eur J Immunol, 31, 812–22.

Wu, X., Briseno, C. G., Durai, V., Albring, J. C., Haldar, M., Bagadia, P., Kim, K. W., Randolph, G. J., Murphy, T. L. & Murphy, K. M. 2016. Mafb lineage tracing to distinguish macrophages from other immune lineages reveals dual identity of Langerhans cells. J Exp Med, 213, 2553–2565.

Yu, Y. A., Hotten, D. F., Malakhau, Y., Volker, E., Ghio, A. J., Noble, P. W., Kraft, M., Hollingsworth, J. W., Gunn, M. D. & Tighe, R. M. 2015. Flow Cytometric Analysis of Myeloid Cells in Human Blood, Bronchoalveolar Lavage, and Lung Tissues. Am J Respir Cell Mol Biol.

Yu, Y. A. & Tighe, R. M. 2018. Isolation and Characterization of Human Lung Myeloid Cells. Methods Mol Biol, 1809, 111–119.

